# A nucleotide code governs Lis1’s ability to relieve dynein autoinhibition

**DOI:** 10.1101/2024.12.30.630615

**Authors:** Indigo C. Geohring, Pengxin Chai, Bharat R. Iyer, William D. Ton, Jun Yang, Amy H. Ide, Sydney C. George, Jaiveer S. Bagri, Samuel V. Baird, Kai Zhang, Steven M. Markus

## Abstract

Dynein-1 is a microtubule motor responsible for the transport of cytoplasmic cargoes. Activation of motility requires it first overcome an autoinhibited state prior to its assembly with dynactin and a cargo adaptor. Studies suggest that Lis1 may relieve dynein’s autoinhibited state. However, evidence for this mechanism is lacking. We first set out to determine the rules governing dynein-Lis1 binding, which reveals that their binding affinity is regulated by the nucleotide-bound states of each of three nucleotide-binding pockets within the dynein motor domain. We also find that distinct nucleotide ‘codes’ coordinate dynein-Lis1 binding stoichiometry by impacting binding affinity at two different sites within the dynein motor domain. Electron microscopy reveals that a 1 Lis1:1 dynein complex directly promotes an open, uninhibited conformational state of dynein, whereas a 2:1 complex resembles the autoinhibited state. Cryo-EM analysis reveals the structural basis for Lis1 opening dynein relies on interactions with the linker domain.

## Introduction

Retrograde transport by the microtubule motor cytoplasmic dynein-1 (hereafter referred to as dynein) supports numerous physiological processes, including tissue morphogenesis, autophagy, and error-free mitosis^10^. This minus end-directed motor complex – which comprises light, light-intermediate, intermediate, and heavy chains – transports numerous cargoes, including membrane-enclosed vesicles, RNAs, proteins, and the nucleus. To engage in processive transport, dynein must first associate with its activating complex dynactin, as well as one of a growing family of cargo adaptor proteins, such as BicD2, Hook3, or Spindly^11, 12^. As a consequence of linking dynein to dynactin and a cargo, these molecules serve to activate dynein motility^13^.

As a check against the untimely or inappropriate delivery of cargoes, dynein and its regulators employ numerous means of regulation that coordinate the initiation of processive motility. Among them are autoinhibitory mechanisms for the 23-subunit dynactin complex (via intra-complex interactions), as well as for several of the adaptor proteins^5, 13, 15-19^. Recent studies have identified that autoinhibition of the adaptors can be relieved via interaction with cargoes^20, 21^, while dynactin can be uninhibited by interactions with dynein itself^5^. For example, contacts between the dynein intermediate chain and the p150 subunit of dynactin can disrupt intra-complex interactions within dynactin that would otherwise maintain it in an autoinhibited state^5^.

In addition to dynactin and adaptors, dynein itself adopts an autoinhibited conformational state referred to as the ‘phi’ particle (due to its resemblance to the Greek letter)^2, 22, 23^. The phi conformation, or analogous states, have been observed for cytoplasmic dynein-1 (from yeast and animals), as well as dynein-2 (responsible for intraflagellar transport)^24, 25^ and the axonemal outer arm dynein^26^. Although somewhat different, these states all involve intra-complex contacts between various dynein complex subunits, and reduce the ability of dynein to bind microtubules, and/or assemble into motile dynein-dynactin-adaptor (DDA) complexes^2, 24, 26^. The importance of the autoinhibited state is apparent from the severe neurodevelopmental diseases that can arise in patients with dynein mutations predicted to disrupt the phi particle^27-29^.

Although relief of autoinhibition is required for assembly of dynein-1 into motile DDA complexes, how it does so is unclear. Recent studies have suggested that the phi particle can be ‘opened’ by the lissencephaly-related protein Lis1^23, 30-33^, a critical conserved regulator that promotes dynein activity by at least several means. In addition to this proposed function, Lis1 also helps link dynein to dynactin during assembly of DDA complexes^5, 30, 31^, and promotes the association of dynein-dynactin complexes with the plus end of dynamic microtubules^34-36^, which is thought to be important for delivery of DD complexes to various cargoes^33, 37-40^. Direct evidence for Lis1 in opening the phi state of dynein, or somehow stabilizing the open state, is currently lacking. This role for Lis1 is based on compelling, but indirect evidence collected from functional (e.g., single molecule motility assays), biochemical (e.g., binding assays), or genetic assays.

The lack of direct evidence to support a role for Lis1 in opening the phi conformation may be due to an incomplete understanding of the ‘rules’ governing dynein-Lis1 binding, and the inability to directly correlate potential structural changes (i.e., phi vs open dynein) with the Lis1-bound state of dynein. One of the complicating factors is the fact that Lis1 has been observed having variable binding modes to dynein: one in which a single WD40 domain from Lis1 is bound to the dynein motor, and another in which two WD40s from a Lis1 dimer is bound to a single motor domain^4, 7, 14, 41^. Although some evidence suggests that the nucleotide-bound state of dynein somehow coordinates Lis1 binding^4, 14, 42, 43^, much remains unknown. For example, it is well established that ATP + vanadate (conditions that mimic the transitional ADP-Pi state^8^) stimulates dynein-Lis1 binding^42^. In addition to these conditions, we recently made the confounding finding that both AMPPNP (a non-hydrolyzable ATP analog) and apo (nucleotide depletion) conditions also promote their binding^4^. These seemingly contradictory results likely stem from the fact that a single dynein motor domain possesses multiple ATP-binding modules^44^ (i.e., the <u>A</u>TPase <u>A</u>ssociated with various cellular Activities, or AAA domains), thus complicating the determination of the exact nucleotide state of the motor domain in previous studies. To understand whether and how Lis1 affects dynein autoinhibition, it is therefore important that we first clarify the rules governing their binding.

In this study we employ a combination of biochemical and electron microscopy-based approaches to understand the rules and the consequences of dynein-Lis1 binding. We find that the nucleotide-bound states of AAA1, AAA3 and AAA4 all coordinate to affect dynein-Lis1 binding affinity. In addition to affecting the strength of binding, we find that specific nucleotide ‘codes’ impact the binding stoichiometry of dynein and Lis1 and determine whether Lis1 binding promotes an ‘open’ state of dynein, or not. We use cryo-electron microscopy to resolve the structural details that account for Lis1 opening phi dynein, which reveals novel dynein-Lis1 contact points, and a structural basis for the distinct binding modes between these two proteins. Our results support a model in which Lis1 can indeed open and/or stabilize an uninhibited conformation of dynein, but only when the AAA domains are in specific nucleotide-bound states.

## Results

### The nucleotide-bound state of AAA1, 3 and 4 all impact dynein-Pac1 binding

We recently found that the microtubule-bound state of dynein impacts dynein-Pac1 (yeast) and dynein-Lis1 (human) binding. Specifically, we found that locking dynein in a <u>m</u>icrotubule-<u>b</u>ound state (MT-B; via protein engineering with a rigid coiled-coil from seryl tRNA synthetase^6^, SRS^CC^; Fig. 1A) significantly reduces dynein-Pac1/Lis1 binding affinity compared with a microtubule-unbound (MT-U) state^4^. Although different nucleotide conditions change the relative binding of Pac1 and Lis1 to both the MT-U and MT-B mutants, the MT-U state exhibits higher affinity for Pac1 in all cases. This led us to ask the following two questions: (1) At which AAA module is each nucleotide condition acting to affect dynein-Pac1 binding? And (2) how does nucleotide occupancy at each AAA domain affect the MT-B/U state to govern dynein-Pac1 binding affinity? To address these questions, we measured the relative ability of full-length Pac1 (a constitutive dimer) to bind monomeric MT-U and MT-B dynein motor domains (dynein_MOTOR_) with mutations that render each AAA module either unable to bind nucleotide (by mutating the Walker A motif, “WA”), or unable to hydrolyze ATP (with Walker B mutations, “WB”; Fig. 1A). We mutated the primary site for ATP hydrolysis (AAA1)^45^, as well as the two other AAA domains that possess ATPase activity: AAA3 and 4^46-49^.

**Figure 1:**
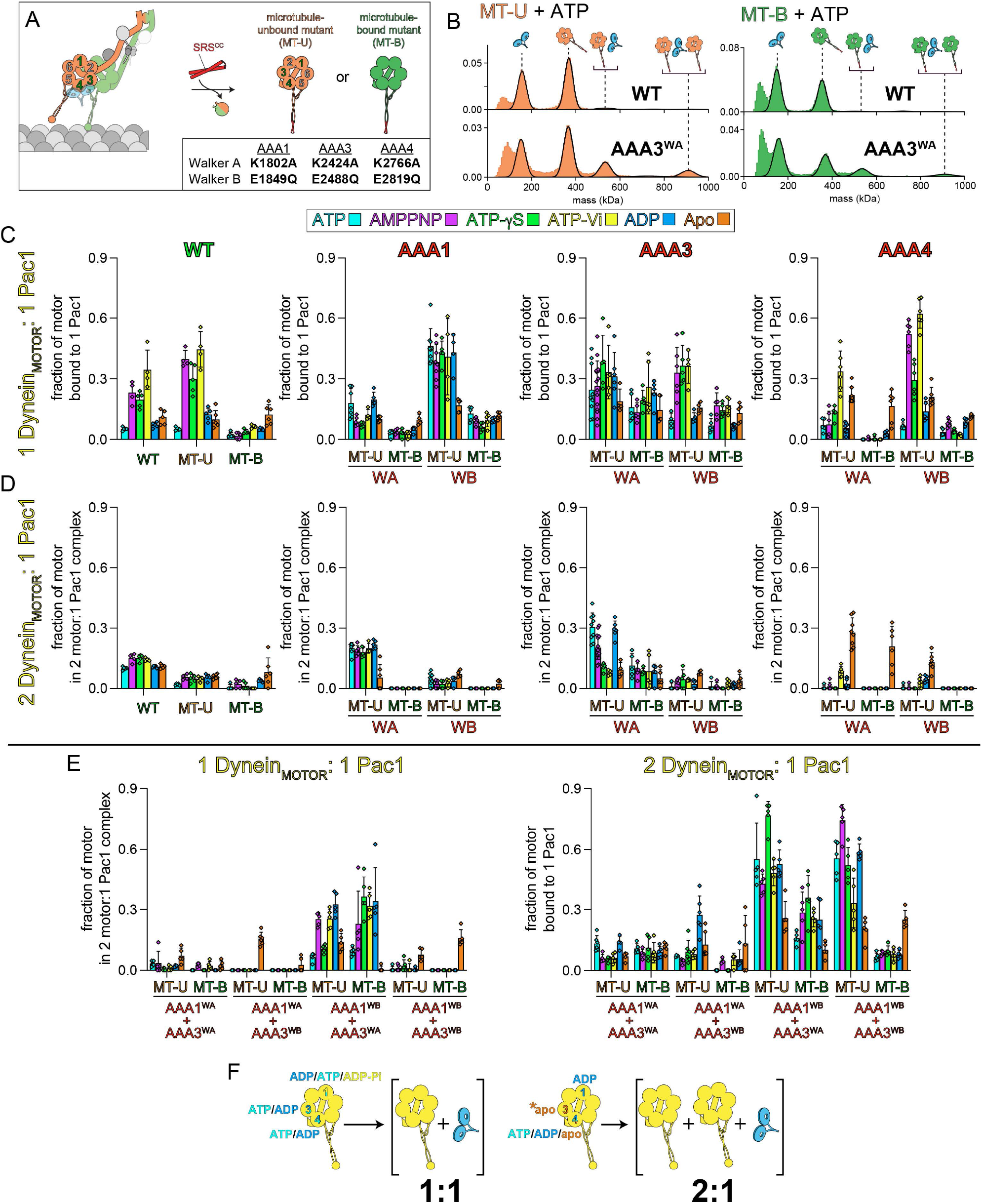
The role of the AAA Walker A and B motifs in dynein-Pac1 binding. **(A)** Cartoon depicting a microtubule-bound full-length dynein complex (with a translucent Pac1 to indicate region of binding) and monomeric dynein_MOTOR_ domains used for binding assays. The fragments were engineered to be locked in either a microtubule-unbound (MT-U) or -bound (MT-B) conformation by replacing the microtubule-binding domain (MTBD) with a rigid coiled-coil derived from seryl tRNA synthetase (SRS)^4, 6^. **(B)** Representative mass histograms obtained from mass photometry-based binding experiments with wild-type and AAA3 Walker A mutant (AAA3^WA^, K2424A) dynein_MOTOR_^MT-U^ and dynein_MOTOR_^MT-B^. **(C - E)** Plots depict results (mean ± standard deviation along with individual data points; each from between 4-15 independent replicates) from mass photometry binding assays. Plots indicate fraction of motors within the 523 kDa peak, which correspond to 1 Pac1^dimer^:1 dynein_MOTOR_ monomer complexes (**C and E, left**), or those within the 897 kDa peak, which correspond to 1 Pac1^dimer^:2 dynein_MOTOR_ complexes (D and E, right; see cartoon schematic above each peak in panel B). **(F)** Results from binding assays indicate nucleotide conditions at each AAA domain that lead to maximal binding of 1:1 or 2:1 complexes. Although a main determinant of 1:1 versus 2:1 binding is the nucleotide state of AAA3 (as indicated by asterisk on “apo-AAA3”), AAA1 also plays a key role.

To measure binding, we mixed equimolar concentrations of dynein_MOTOR_ and Pac1 in the absence or presence of various nucleotides, and quantitatively assessed the relative proportion of unbound and bound species via mass photometry. As we noted previously^4^, two distinct complexes become apparent: a 1:1 dynein_MOTOR_:Pac1^dimer^ complex (expected mass = 513 kDa), and a second species that is indicative of a 2:1 dynein_MOTOR_:Pac1^dimer^ complex (expected mass = 872 kDa; Fig. 1B)^4^.

Our results (Fig. 1C and D) show that both nucleotide-bound and -unbound (apo) AAA1 and AAA3 pockets can stimulate Pac1 binding, albeit to different extents, and in different binding ratios. Specifically, whereas the AAA1^WB^ and AAA3^WB^ mutants form 1:1 complexes with Pac1 to relatively high degrees, the corresponding WA mutants bind Pac1 in 2:1 ratios. The results from the various nucleotide treatments further indicate that both apo-AAA1 and apo-AAA3 must be combined with at least one other nucleotide-bound AAA module for maximal 2:1 Pac1 binding, as depletion of nucleotide reduced complex formation for each (*e*.*g*., see apo conditions for the AAA1^WA^ and AAA3^WA^ mutants).

Our findings with the AAA4^WA^ and AAA4^WB^ mutants show that they exhibit a similar pattern of binding compared to each other and when compared to the wild-type motor, except that the AAA4^WA^ mutant binds Pac1 to lower extents for most conditions, while the AAA4^WB^ mutant binds to greater extents. These results indicate that the AAA4 pocket must be bound to nucleotide for dynein to adopt a high Pac1 affinity state.

We next assessed the combined effect of the AAA1 and AAA3 mutants. Although the AAA1^WA^ mutant alone assembles the 2:1 complex to a high degree, pairing this mutation with either AAA3^WA^ or AAA3^WB^ led to poor Pac1 binding (Fig. 1E). On the other hand, we found that even higher degrees of complex formation can be achieved when we combined AAA1^WB^ with either AAA3^WA^ or AAA3^WB^. Notably, whereas the former efficiently assembles the 2:1 complex with Pac1, the latter does so to a very low degree. Although these data suggest that AAA3 is the main determinant of binding stoichiometry, we also note one instance in which AAA1 can also do so. Specifically, when the AAA3^WA^ single mutant is treated with ATP + vanadate (Vi), we observe low degrees of 2:1 complex assembly, but high levels of 1:1 binding. Given AAA1 is the likely site at which ADP-Vi is bound^4, 8^, these data indicate that AAA1 and AAA3 can both affect the affinity and stoichiometry of dynein-Pac1 binding.

In most cases with the AAA1 and AAA3 mutants (with apo conditions being the main exception), the microtubule-bound and -unbound states behave as we noted previously with the wild-type proteins. Specifically, the MT-U state binds to Pac1 to a higher degree than the corresponding MT-B state, which is consistent with the model that microtubule-binding by dynein reduces dynein-Pac1/Lis1 binding affinity^4^. This is true for both the 1:1 and the 2:1 complexes, indicating that the interaction surfaces that account for these distinct species are both subject to microtubule-binding-induced conformational changes, and that the MT-U/MT-B state is largely dominant to the nucleotide-bound state.

### A Pac1 dimer bridges two monomeric motor domains

We next sought to determine the nature of the 2 dynein_MOTOR_:1 Pac1^dimer^ complex. To this end, we enriched for this species of Pac1-bound AAA3^WA^ dynein_MOTOR_^MT-U^ mutants (assembled in the presence of ADP) using size exclusion chromatography, and assessed their morphology via single particle negative stain EM. This revealed the presence of pairs of motor domains situated in close apposition to each other (Fig. 2A). 2D classification analysis reveals that the motor domain pairs are sufficiently homogeneous that several of the classes reveal detailed images of each motor domain within a pair (Fig. 2B). Thus, the motor pairs are held together with fairly consistent spacing and conformational conformity with respect to each other. We also note that the Pac1^dimer^-linked motors are able to freely rotate with respect to one another and exist in a roughly equal mixture of “aligned” (with the coiled-coil “stalk” domains pointing in the same direction) and “inverse” configurations (with stalks pointing in opposite directions; Fig. 2C). The quality of our negative stain EM averages permitted us to manually fit a previously obtained 3D model (of a Pac1-bound monomeric motor domain^7^) to them with a high degree of confidence, and to unambiguously identify densities corresponding to the motor domains as well as the bound Pac1. We then manually docked various elements from previous high-resolution structures, including the dynein-2 motor domain (PDB 4RH7^8^), WD40 domains from Lis1 (PDB 8FDT^4^), and the N-terminal LisH domain (PDB 1UUJ^50^), which verified the rotational freedom for the Pac1-bound motor domains.

**Figure 2:**
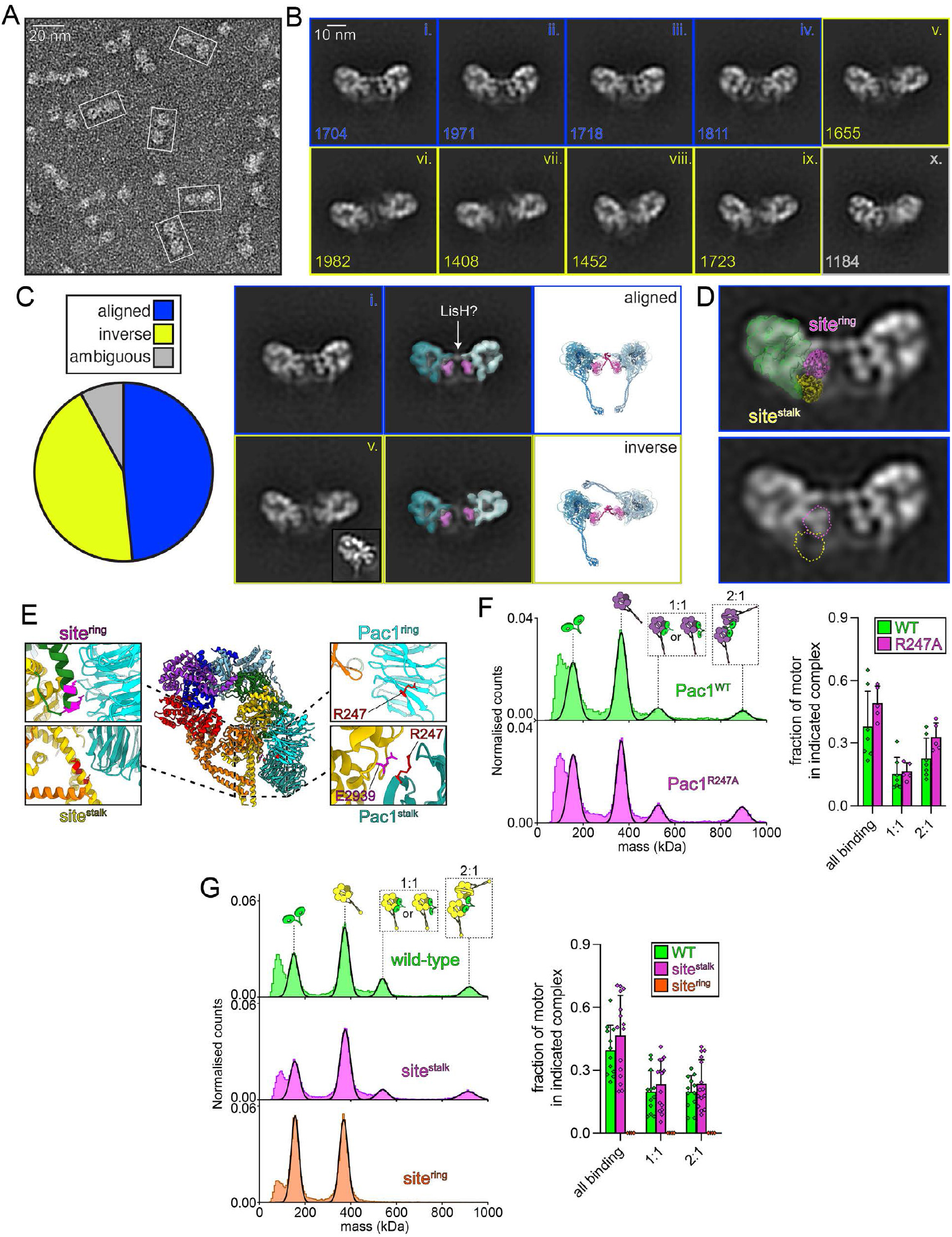
Pac1 bridges two dynein_MOTOR_ domains via site^ring^ contacts. **(A and B)** Representative electron micrographs **(A)** and 2D class averages **(B)** from negative stain grids prepared with Pac1-bound dynein_MOTOR_^MT-U/K2424A^ (in the presence of ADP). White boxes in panel **A** delineate closely apposed pairs of motor domains. Classes i-iv represent motor domains aligned in a parallel manner, classes v-ix are in an inverse orientation, and class x is ambiguous. **(C)** Quantitation of indicated motor orientations from 2D classification along with superimposition of 3D maps from EMD-6008^7^ (one on each motor domain; middle panel) on classes i and v from panel **B**. Colors depict motor domain (shades of cyan) and Pac1 (magenta). Various elements from high-resolution structures were manually docked into the EM density (motor domain from 4RH7^8^, linker from 3VKG^9^, and WD40 domain from 8FDT^4^). Although unclear, the unassigned density may represent the N-terminal Lis1H dimerization domain of Pac1. Inset shows focused 2D classification of a monomeric motor domain. **(D)** 2D class average (class i) with superimposition of 3D map from EMD-8706^14^, which represents a dynein motor domain with Pac1 WD40 domains bound at both site^ring^ (magenta) and site^stalk^ (yellow). Note the lack of EM density for Pac1 at site^stalk^ in our 2D averages (dashed yellow outline). **(E)** Close-up view of site^ring^ and site^stalk^ in yeast dynein_MOTOR_ (with a AAA3^WB^ mutation)^14^. **(F and G)** Representative mass photometry histograms (left) and plots (mean ± SD; right) showing binding between dynein_MOTOR_^MT-U/K2424A^ and Pac1^R247A^, **(F)** or between Pac1 and either WT, site^ring^ (K2721A, D2725G, E2726S, and E2727G) or site^stalk^ (E3012A Q3014A N3018A) mutants of dynein_MOTOR_^M2424A^ (**G**; each from between 5-15 independent replicates).

Each Lis1 and Pac1 WD40 domain within a dimer binds to two distinct sites on a single dynein motor domain: one between AAA3 and 4 (site^ring^)^7, 51^, and another at the base of the coiled-coil stalk (site^stalk^)^14, 43^. The close alignment of our 2D averages to that from the 3D model (EMD-6008^7^) strongly suggests that Pac1 is likely bound to site^ring^. Moreover, comparison of our 2D averages to a recent structure of dynein bound to two Lis1 WD40 domains (one at each site)^14^ reveals a lack of density at site^stalk^ (Fig. 2D). To confirm this contact point we generated three point mutants: (1) dynein_MOTOR_ fragments with mutations at site^stalk^ or (2) site^ring^ that perturb Pac1 binding^14, 51^, and (3) Pac1 with a mutation predicted to disrupt contacts at site^stalk^, but not site^ring^ (R247A; Fig. 2E). For experiments 1 and 2, we used dynein_MOTOR_^K2424A^ fragments with the native wild-type stalk and microtubule-binding domain, whereas we used dynein_MOTOR_^MT-U/K2424A^ for the latter. These experiments reveal that disruption of neither site^stalk^ nor the Pac1 R247A mutation affects assembly of the 2:1 complex (Fig. 2F and G). However, mutation of site^ring^ eliminates this species, indicating that site^ring^, but not site^stalk^, is indeed required for 2:1 binding.

### Binding modes are governed by differential Pac1 affinity at two distinct sites

Our data thus far suggest the possibility that the two different dynein-Pac1 complexes are a consequence of the two Pac1 binding sites on dynein separately modulating their affinity. In particular, we hypothesize that an apo-AAA3 in combination with an ADP-AAA1 simultaneously leads to a higher binding affinity for Pac1 at site^ring^ and a lower affinity at site^stalk^. An ATP-AAA3 with either an ATP or ADP-Pi-bound AAA1 on the other hand leads to weakened Pac1 affinity at site^ring^, and increased affinity at site^stalk^.

To test whether site^ring^ changes affinity due to the AAA3 state, we assessed the ability of monomeric Pac1 WD40 domains (*i*.*e*., Pac1^ΔN^, which lacks the N-terminal dimerization domain) to bind to dynein_MOTOR_. Previous studies have shown that this fragment binds to site^ring^, and not site^stalk^, and thus can be used to interrogate binding affinity at this site^4, 7^. We noted very little binding between wild-type dynein_MOTOR_ and Pac1^ΔN^, except in ATP-Vi and apo conditions (Fig. 3A), indicating this fragment binds dynein with low affinity (likely due to lack of avidity). However, we observed robust 1:1 complex formation for the AAA3^WA^ mutant in all conditions except apo, indicative of an apo-AAA3 promoting high Pac1binding affinity at site^ring^. If instead of enzymatically depleting nucleotide, we simply reduce ATP concentrations via dilution in nucleotide-free buffer (to 10 µM), we observe robust binding for the AAA3^WA^ mutant. The same is true for the wild-type dynein motor. This is consistent with AAA1 (and AAA4) requiring nucleotide for high affinity binding at site^ring^. In contrast to the AAA3^WA^ mutant, we observe no binding between the AAA3^WB^ mutant and Pac1^ΔN^ in all conditions, consistent with ATP-AAA3 causing lower affinity at site^ring^.

**Figure 3:**
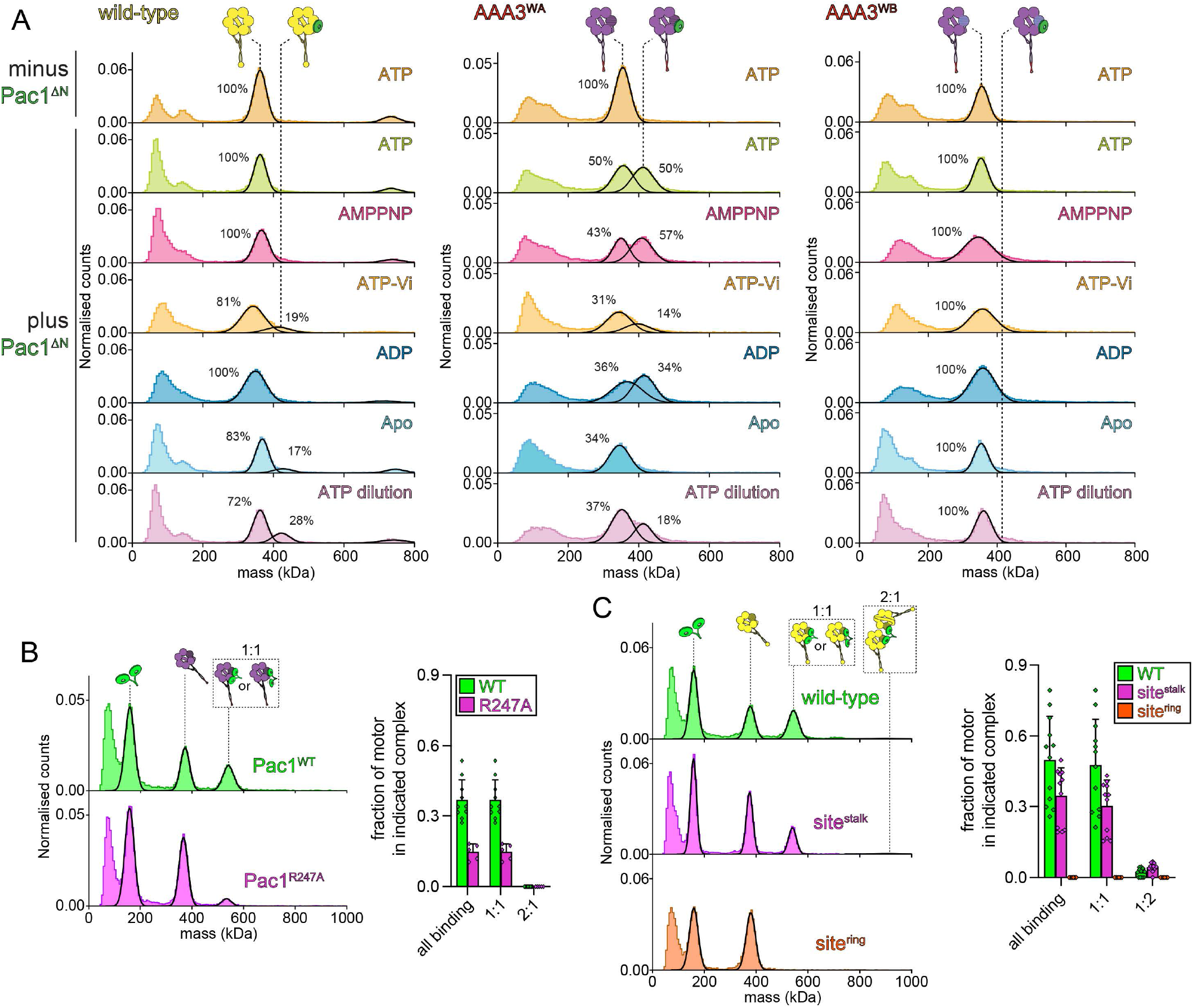
AAA1 and AAA3 coordinate Pac1-binding affinity at both site^ring^ and site^stalk^. **(A)** Representative mass photometry histograms showing binding between monomeric Pac1 (Pac1^ΔN^; Δ1-143) and indicated monomeric dynein_MOTOR_ (WT, left; MT-U/K2424A, middle; MT-U/E2488Q). Each are representative of at least 3 independent replicates. Note that the relatively low mass of unbound Pac1^ΔN^ makes it undetectable by mass photometry in our conditions. **(B and C)** Representative mass photometry histograms (left) and plots (mean ± SD, along with all data points; right) showing binding between full-length Pac1 (WT or R247A) and either dynein_MOTOR_^MT-U/E2488Q^ **(B)** or dynein_MOTOR_^E2488Q^ (**C**; each from between 5-12 independent replicates).

To determine whether Pac1 affinity at site^stalk^ is affected by the nucleotide-bound state of AAA3, we tested the importance of this site via mutagenesis. Consistent with the importance of site^stalk^ in dynein-Pac1 binding when AAA3 is bound to ATP, we observe a significant reduction in binding between Pac1^R247A^ and the AAA3^WB^ mutant (Fig. 3B). We observe a similar reduction in binding between wild-type Pac1 and the AAA3^WB^ dynein mutated at site^stalk^ (Fig. 3C). These data are also supported by previous cryo-EM data that show that a site^stalk^-bound Pac1 WD40 domain only becomes apparent when AAA3 is bound to ATP (via an AAA3^WB^ mutation)^14^. Taken together, our data support a model in which both site^ring^ and site^stalk^ affinity are modulated by the nucleotide-bound states of AAA1 and AAA3.

### Nucleotide-bound state of dynein dimers coordinates Pac1binding stoichiometry

Given a dynein complex consists of two motor domains with as many as four Pac1 WD40 domain binding sites (two site^ring^ and two site^stalk^), we wondered how the different dynein-Pac1 binding modes described above translate to such a context. To this end, we purified a well-characterized glutathione S-transferase (GST)-dimerized dynein_MOTOR_ fragment with or without AAA3^WA^ or AAA3^WB^ mutations, and assessed their binding to Pac1 in different nucleotide conditions (Fig. S1). In contrast to the monomeric dynein_MOTOR_ fragments, we observed no evidence for Pac1 dimers linking two distinct dimeric GST-dynein_MOTOR_ fragments in any conditions (note the lack of an 1844 kDa species), including those that enrich for 2 dynein_MOTOR_:1 Pac1^dimer^ species (*i*.*e*., AAA3^WA^ mutant with ATP or ADP). We ensured this wasn’t the case over a range of concentrations, even when Pac1 is present at 3X molar excess. Instead, our data reveal that each motor dimer bound to either one Pac1 dimer (1:1 ratio), or two (1:2 ratio), depending on the conditions, which we describe below.

Whereas the wild-type and AAA3^WB^ GST-dynein_MOTOR_ assemble into 1:1 and 1:2 complexes – both in conditions that promote high Pac1 affinity (AMPPNP and ATP-Vi) – the AAA3^WA^ mutant forms predominantly 1:1 complexes with Pac1. As expected from our data above, the AAA3^WA^ mutant bound Pac1 best in the presence of ATP, AMPPNP, and ADP, likely due to the adoption of a high affinity state at site^ring^. In all these cases, we observed an almost exclusive 1:1 stoichiometry of binding. This is consistent with each WD40 domain of a Pac1 dimer binding to site^ring^, with likely no binding at site^stalk^, likely due to this latter site adopting a low affinity state. Of note, the AAA3^WA^ mutant can assemble a 1:2 complex with Pac1 in the presence of ATP-Vi, conditions that promote strong 1:1 binding with the monomeric dynein motor domain, but very little 2:1 binding (Fig. 1), which is indicative of site^ring^ and site^stalk^ adopting low and high affinity states, respectively. These data indicate that an apo-AAA3 combined with an ATP- or ADP-AAA1 leads to assembly of 1:1 dynein-Pac1 complexes, whereas ATP-AAA3 or ADP-Pi-AAA1 promotes 1:2 stoichiometry.

To determine whether these findings apply to the native dynein complex, we purified full-length yeast dynein complexes (wild-type, AAA3^WA^, and AAA3^WB^) from insect (Sf9) cells. Mass photometry of the purified complexes reveal masses of 1255 ± 13 kDa, 1257 ± 11 kDa, and 1257 ± 10 kDa for the WT, AAA3^WA^, and AAA3^WB^ mutants, respectively, which very closely match the predicted mass of these complexes (1217 kDa). Pac1 binding experiments with each reveal an identical nucleotide-dependent pattern for the fulllength dynein to that of GST-dynein_MOTOR_ (Fig. 4). Specifically, AMPPNP and ATP-Vi stimulate assembly of both 1:1 and 1:2 species with the wild-type and AAA3^WB^ mutant, whereas the AAA3^WA^ mutant assembles into 1:1 complexes in the presence of ATP and ADP, and 1:2 complexes in ATP-Vi conditions. Taken together, we identify a nucleotide-dependent code that coordinates the assembly of two distinct dynein-Pac1 complexes.

**Figure 4:**
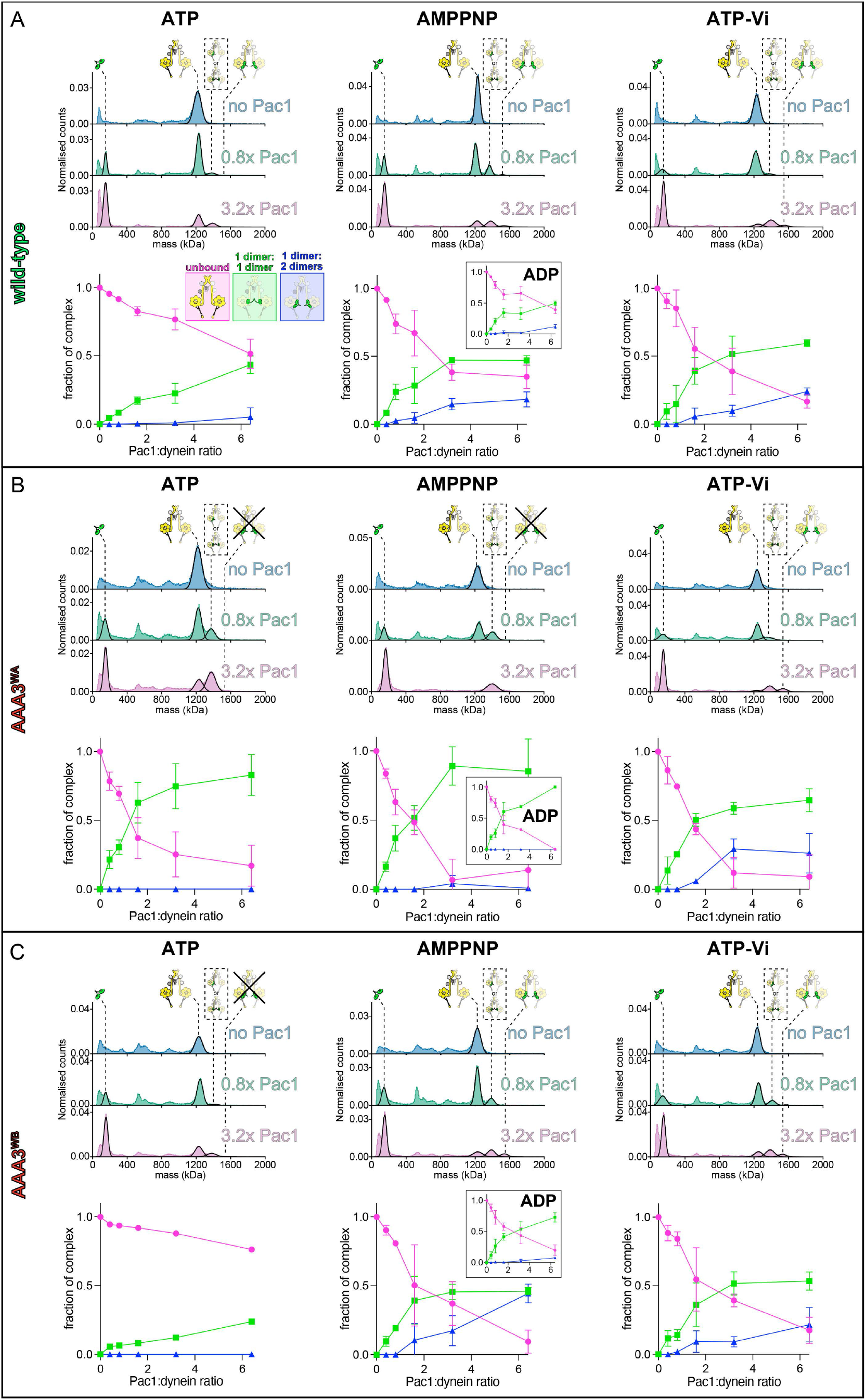
Nucleotide occupancy coordinates stoichiometry of binding between Pac1 and full-length dynein. Representative mass photometry histograms and associated plots (mean fraction of binding ± SD; each from 3-6 independent replicates) depicting extent and stoichiometry of binding between Pac1 and indicated full-length dynein complex (purified from SF9 cells) at a range of dynein:Pac1 ratios.

### The ability of Pac1 to ‘open’ dynein is determined by their binding stoichiometry

In light of our identification of two distinct dynein-Pac1 complexes, and the conditions of their assembly, we next sought to determine how they each impact dynein function. Given the proposed but untested model that Pac1 releases dynein autoinhibition, we focus on how these distinct Pac1binding modes might influence dynein’s adoption of the autoinhibited ‘phi’ particle. To this end, we mixed full-length AAA3^WA^ and AAA3^WB^ dynein mutants with Pac1 in conditions that favor either 1:1 (AAA3^WA^ with ATP) or 1:2 (AAA3^WB^ with AMPPNP) binding stoichiometry and assessed dynein morphology by negative stain EM. In order to enable unambiguous analysis of the resulting dynein-Pac1 complexes, we used sufficiently high Pac1:dynein ratios such that dynein binding was close to saturation (Fig. 5A and B).

**Figure 5:**
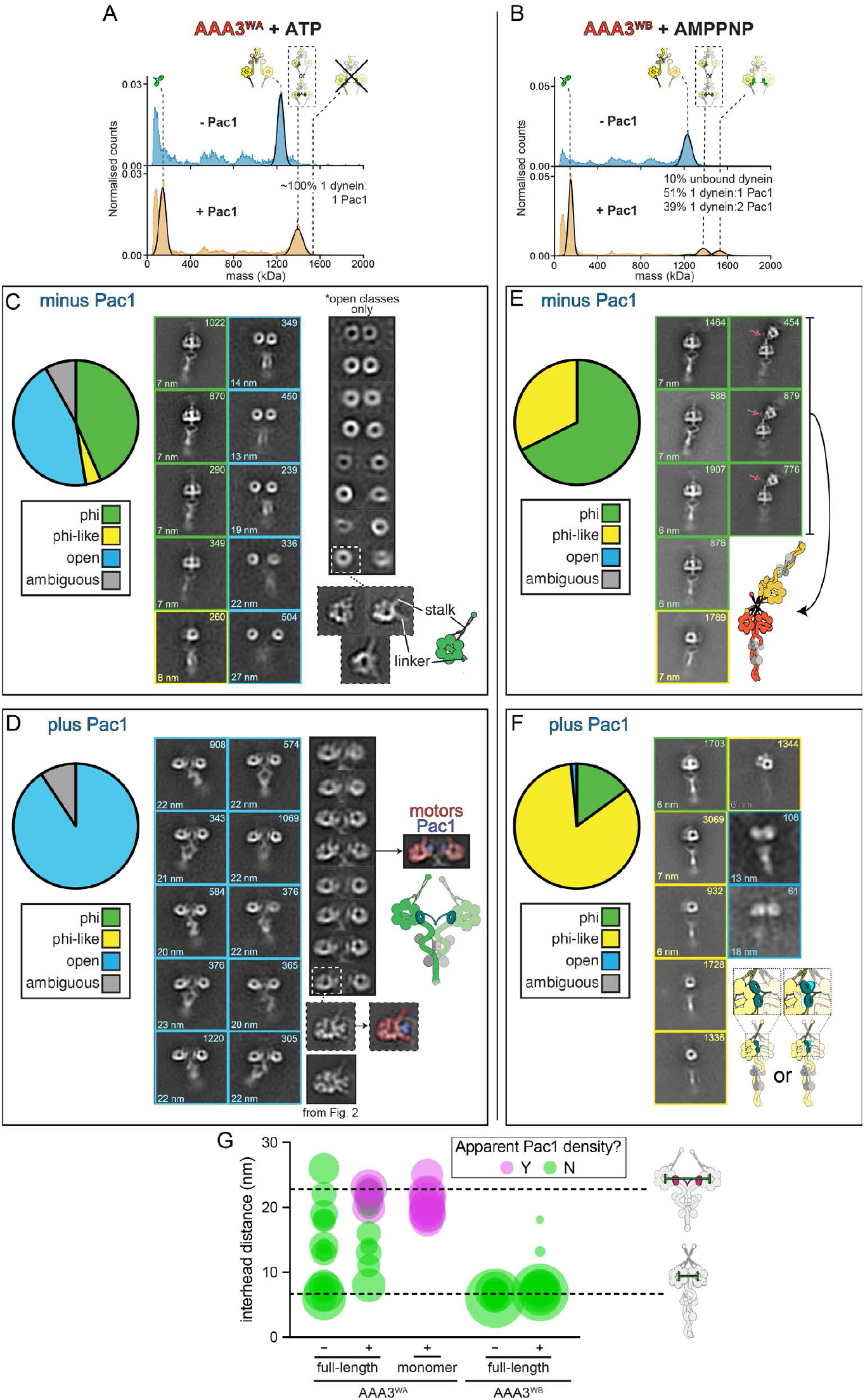
Nucleotide occupancy and Pac1 binding mode dictate conformation of full-length dynein. **(A and B)** Mass photometry histograms of samples containing full-length dynein (AAA3^WA^ or AAA3^WB^ mutant) with or without excess Pac1 that were used to directly prepare negative stain EM grids, images of which are shown in panels **C – F. (C – F)** Representative 2D class averages showing indicated full-length dynein (AAA3^WA^ or AAA3^WB^ mutant) in the absence **(C and E)** or presence **(D and F)** of Pac1. Pie graphs show relative fraction of indicated conformational state (determined from 2D averages). Numbers indicate the number of particles in each class (top) and the measured distance between the motor domains (from center-to-center, bottom; note those missing measurements are due to inability to clearly identify motor domain centers). For panels **C and D**, focused classifications of motor dimers or monomers are shown (a Pac1-bound monomer from Figure 2C is shown for comparison in panel **D**). The poor image quality for the dimers in panel **C**, but not the monomers, indicate a large range of conformational heterogeneity within the motor dimer pairs. Note the improved resolution for motor dimer averages in panel **D**, indicating the reduced heterogeneity for Pac1-bound motor dimers compared to those unbound from Pac1 (Pac1 and dynein motor domain pseudo-colored in blue and red, respectively). We observed a novel species in which two phi dynein dimers are bound via apparent contacts between the motor domains (see cartoon depiction in panel **E**) that appear to stabilize the stalk-MTBD region (red arrows indicate improved resolution of this region). **(C)** Bubble plot of interhead measurements taken from 2D class averages for indicated species. Bubble size reflects proportion of particles per 2D class, whereas green/magenta indicate whether Pac1 density can be discerned from each average. Monomer data was measured from those images shown in **Figure 2C**. Dashed lines indicate mean interhead distance for Pac1-bound AAA3^WA^ mutant (22 nm) and phi dynein (6.9 nm). Each dataset is representative of 2 independent replicates.

Assessment of the AAA3^WA^ mutant in the absence of Pac1 reveals that approximately half of the dyneins adopt an autoinhibited state, with the remainder existing in an open state (Fig. 5C). 2D class averages of phi dynein reveal an overall morphology that is indistinguishable from wild-type dynein complexes (Fig. S2). Focused classification of the motor pairs in the open configuration reveals very low-resolution pictures of the motor domains that adopt a wide range of inter-head spacing (Fig. 5G). The low resolution is a consequence of the high degree of flexibility for the open dyneins, since focused classification of individual motor domains from the same data set reveal sufficient detail to identify various structural features (*e*.*g*., the coiled-coil stalk and linker; Fig 5C, bottom).

Addition of Pac1 to the AAA3^WA^ mutant results in a striking transition of dynein to an open state (Fig. 5D). Although we observe a small fraction of dyneins that closely resemble the phi particle (“phi-like”), the large majority of dyneins appear to be in an open conformation in the presence of Pac1. Unlike the open state for Pac1-unbound dynein, those bound to Pac1 adopt a more consistent morphology with less rotational freedom, and a smaller range of spacing between the motor domains, as evidenced by the higher resolution 2D averages for the full-length complex, as well as our measurements of inter-head spacing (Fig. 5G). Note that the interhead spacing with the Pac1-bound motor pairs is very similar to that of the 2 dynein_MOTOR_ : 1 Pac1^dimer^ complexes (Fig. 2B and 5G). Focused classification of the motor pairs permits identification of densities that correspond to Pac1 WD40 domains bound to site^ring^ on each motor domain. This was even more apparent from focused classification of the individual motor domains, which reveal a 2D average that is almost identical to that obtained from the monomeric Pac1-bound motor domains (see Fig. 2). These observations indicate that Pac1 indeed has the capacity to stabilize and/or promote an open conformation of the full-length dynein complex, at least when AAA3 is in an apo state and AAA1 is bound to either ADP or ATP. They also show that Pac1 links the motor domains together in a manner that restricts their motion with respect to each other.

A similar analysis of the full-length AAA3^WB^ dynein mutant (in the presence of AMPPNP) reveals that it has a high propensity to adopt the autoinhibited conformation in the absence of Pac1 (Fig. 5E). In fact, unlike wild-type and the AAA3^WA^ mutant, we observed no apparent classes for open dynein with this mutant. In addition to the phi particle, we observed a novel species in which two phi dyneins are bound via apparent contacts between the coiled-coil stalks. The resolution of this species, even by negative stain EM, is sufficient to conclude that this particle is morphologically homogeneous, and likely represents a stably bound dimer of phi dynein dimers. Of particular note, the coiled-coil stalk and microtubule-binding domains, which are normally too flexible to be visible in 2D averages (either by negative stain, or cryo-EM) is clearly apparent in the 2x-phi dynein species, suggesting the inter-dimer contact stabilizes this region (Fig. 5E; see red arrow).

In contrast to the AAA3^WA^ mutant, addition of Pac1 to the AAA3^WB^ mutant does not result in dynein adopting an open conformation (Fig. 5F). Moreover, 2D averages do not reveal a large proportion of canonical phi dynein species. Instead, the majority of dyneins in these conditions are ‘phi-like’, which have characteristics of phi dynein, but are clearly distinct. The low resolution of the 2D averages of these species suggest a potential lack of symmetry in this phi-like state, and may indicate that Pac1 binding partly opens phi dynein, but only to a minor extent. Thus, whereas the 1:1 dynein-Pac1 complex adopts an open state, the 2:1 complex does not.

### Structural basis for 1 dynein:1 Pac1 complex assembly

To gain insight into the mechanism by which dynein binds Pac1 in a 1:1 ratio, thereby opening phi dynein, we sought to obtain a representative cryo-EM structure. To this end, we mixed full-length wild-type dynein with or without Pac1 in the presence of low concentrations of ATP (0.1 mM; Figs. 6A and S3). For the purpose of this study, we focus exclusively on the motor domains of open dyneins to understand the nature and consequence of dynein-Pac1 binding.

**Figure 6:**
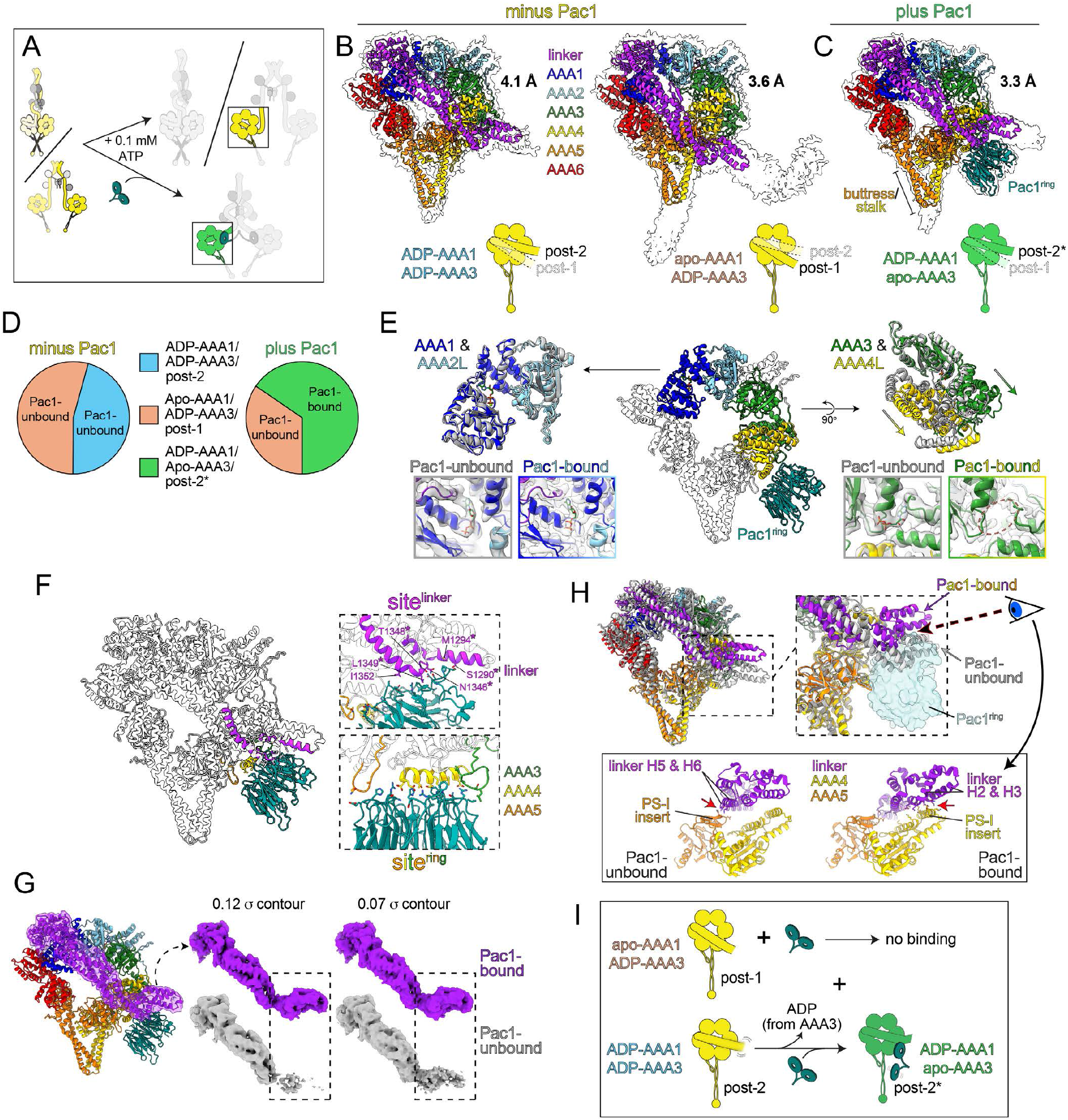
Cryo-electron microscopy of full-length dynein in the absence or presence of Pac1. **(A)** Schematic of experimental setup. Full-length wild-type dynein was incubated with or without Pac1 in the presence of 0.1 mM ATP prior to preparing grids for freezing. We performed a focused classification of individual motor domains of open dyneins. (B) Molecular models of dynein motor domains without **(B)** or with Pac1 **(C)** along with corresponding density maps (indicated with outlines). Resolution of each structure is indicated. Subdomains are color-coded as indicated. Nucleotide occupancy at AAA1 and AAA3 and linker position are indicated by cartoon below each (post-2* indicates a ‘modified post-2’ state; see text). **(D)** Pie graphs showing relative fraction of dynein motors with indicated linker position and nucleotide occupancy at AAA1 and AAA3 for those motors without and with Pac1. Note that the post-2* state, and the presence of an apo-AAA3 are both unique to those bound to Pac1. **(E)** Close-up views comparing AAA1 and AAA3 pockets in the absence (gray) or presence (blue and cyan for AAA1 and AAA2L; green and yellow for AAA3 and AAA4L) of Pac1. Note for minus Pac1, we used the model with ADP-AAA1/ADP-AAA3 and post-2 linker. Arrows in right images depict opening of AAA3 pocket in the presence of Pac1. Insets depict molecular models with EM density overlaid. Red dashed line circle delineates empty AAA3 pocket. **(F)** Close-up view of molecular model showing Pac1-dynein motor contacts at site^ring^ and new site^linker^. Residues marked with asterisks are those mutated in site^linker^ mutant (see Fig. 7 and text). **(G)** Molecular model and EM densities with two different contour settings illustrating that a Pac1 bound to dynein reduces flexibility of the N-terminal region of the linker (see regions within dashed box). **(H)** Comparison of Pac1-bound and unbound (the latter with ADP-AAA1/ADP-AAA3 and post-2 linker) dynein motors reveal a shift of the linker N-terminus away from the stalk and microtubule-binding domains. Close-up views show the linker in two slightly different post-2 docking modes: one in which the linker contacts residues in the PS-I (pre-sensor-I) of AAA5 (minus Pac1), and the other in which the linker contacts residues in the AAA4 PS-I (plus Pac1). **(I)** Model indicating that high affinity binding of Pac1 to site^ring^ and site^linker^ requires ADP binding to AAA1, and ADP release from AAA3.

In the absence of Pac1, we obtained two classes: one possesses ADP-AAA1 and ADP-AAA3, and another with an apo-AAA1 and ADP-AAA3 (Figs. 6B and S4). Whereas those motor domains with an apo-AAA1 exhibit a canonical post-powerstroke linker (“post-1”; docked at AAA5), those with an ADP-AAA1 have the linker in the recently identified post-2 state, in which the linker is docked at AAA4^52^. A post-2 docking mode, which has not yet been observed with yeast dynein, was also recently identified in human dynein-1 and axonemal outer arm dynein^1, 52^, indicating this may be a crucial aspect of the mechanochemical cycle for all dyneins.

In the presence of Pac1, we observe only a single class of dynein motor-bound Pac1, in which AAA1 (and AAA4) is bound to ADP, and AAA3 is in an apo state (Figs. 6C, S3 and S4). These dyneins are bound to a single Pac1 WD40 domain via site^ring^. This observation aligns with our biochemical data indicating that a single WD40 domain binds at site^ring^ when AAA1 is bound to ADP and AAA3 is in an apo state. Unlike previous cryo-EM structures of Pac1- or Lis1-bound dynein motor domains, our Pac1-bound dynein structure possesses a linker that is in a post-powerstroke state in which it is docked at AAA4 in the post-2 state. In addition, the AAA1 pocket is in an open state, and the motor appears to be in a high microtubule-binding affinity state based on the conformation of the buttress and stalk (Fig. 6C). Note this structure resembles a previously obtained low-resolution map (∼20 Å) obtained for a monomeric dynein_MOTOR_ bound to Pac1^7^, and also a previously determined cryo-EM structure solved for Pac1 bound to a monomeric dynein_MOTOR_ with a truncated linker^14^ (5VH9, 7.7 Å).

Comparison of Pac1-bound and unbound dynein reveals that Pac1 shifts the equilibrium of full-length dynein conformations (Fig. 6D). Although we observe motor domains with a post-2 linker in the absence and presence of Pac1, an apo-AAA3 was exclusive to those bound to Pac1. This indicates that Pac1 binding to dynein stabilizes and/or promotes ADP release from AAA3. Comparison of the AAA3 nucleotide binding pocket reveals that the Pac1-bound motor possesses a more open AAA3 pocket that is reflective of its empty state (Fig. 6E, right, and Fig. S4B). Close inspection of the Pac1-bound dynein structure reveals previously identified contacts at site^ring^, but also a novel interaction between Pac1 and the post-2 state linker. Although in both Pac1-unbound and -bound states the linkers are in post-2 positions, there are two apparent differences. First, the linker-Pac1 contact (at site^linker^; Fig. 6F) appears to stabilize the N-terminal region of the linker (Fig. 6G) in a position that is more distal from the stalk (Fig. 6H). This N-terminal region is often truncated in previous structures. Second, Pac1 binding shifts the linker from contacting AAA5 (via the PS-I insert) to instead contact the PS-I insert at AAA4 (Fig. 6H, bottom). Taken together, these data indicate that Pac1 binding stabilizes the linker in a modified post-2 conformation (see “post-2*” in Fig. 6C and D), and the AAA3 pocket in an apo state (Fig. 6I). In light of the fact that the AAA3^WA^ MT-B mutant exhibits lower affinity for Pac1 than the AAA3^WA^ MT-U (Fig. 1D), we posit that microtubule-binding by dynein likely promotes dissociation of this complex, thus permitting ATP binding by AAA3, and subsequent motility.

### Pac1-linker contacts account for different binding modes

Although the Pac1-site^ring^ contacts in our structure largely overlap with those observed in previously obtained dynein_MOTOR_-Pac1 and Lis1 structures^3-5, 7, 14, 31, 41^, a close inspection reveals small but significant differences. We compared our structure to those with two apparent WD40s bound to the motor domain: one at site^ring^ and another at site^stalk^. Notably, this includes a structure solved for an AAA3^WB^ mutant (mixed with ATP-Vi)^14^ that binds to two WD40 domains as a consequence. This comparison reveals that Pac1 is shifted downward towards site^stalk^ in our apo-AAA3 structure (by 7.6 Å when compared to a yeast dynein_MOTOR_-Pac1 structure, 7MGM; Fig. 7A and B). This downward shift, which may be due to the post-2* linker-Pac1 contacts, appears to occlude binding of a second WD40 domain at site^stalk^ (Fig. 7B; note steric clashes become apparent upon docking a Pac1 WD40 at site^stalk^). These observations suggest that the linker-Pac1 contacts may account for both (1) the increase in affinity of Pac1 for site^ring^ (by providing additional contacts at site^linker^), and (2) the decrease in affinity of Pac1 for site^stalk^ (due to steric occlusion).

**Figure 7:**
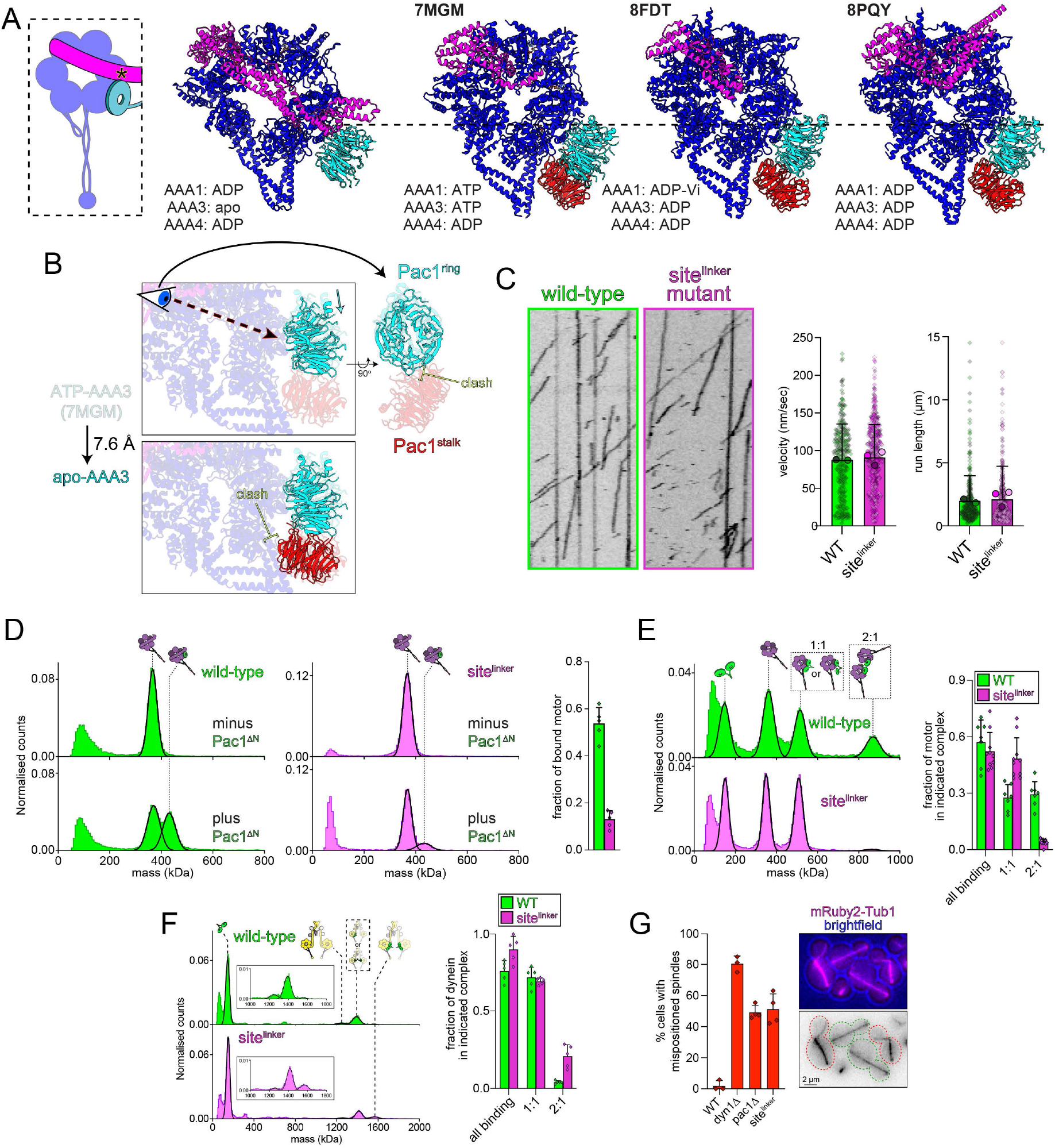
Pac1-site^linker^ contacts account for different dynein-Pac1 binding modes. **(A)** Cartoon and various structural models comparing the position of the site^ring^-bound Pac1 between the various structures with two WD40 domains bound (left-to-right: this study, 7MGM, 8FDT, and 8PQY^3-5^). Asterisk in cartoon delineates approximate region of site^linker^ contacts mutated in subsequent panels. Nucleotide occupancy for AAA1, 3 and 4 are indicated. **(B)** Comparison of site^ring^-bound Pac1 between our structure and a yeast dynein with two WD40s bound (7MGM)^3^. Note the downward shift of Pac1 is predicted to occlude a site^stalk^-bound Pac1 due to clashes at this site. **(C)** Representative kymographs and quantitation of single molecule motility assays performed with either wild-type or site^linker^ mutant variants of GST-dynein_MOTOR_. Small diamonds, all data points; larger circles, means for independent replicates; bars, mean ± standard deviation (for WT and site^linker^, n = 309 and 473 motors from 2 and 3 independent replicates, respectively). **(D and E)** Representative mass photometry histograms showing wild-type or site^linker^ mutant monomeric dynein_MOTOR_^MT-U/K2424A^ in the absence or presence of either monomeric (Pac1^ΔN^; **D**) or dimeric (**E**) Pac1. Plots show mean ± SD along with independent replicates. **(F)** Representative mass photometry histograms and associated plots (mean fraction of binding ± SD) depicting extent and stoichiometry of binding between Pac1 and either full-length dynein^K2424A^, or dynein^K2424A^ with site^linker^ mutations (in the presence of AMPPNP). For panels D – F, n = 5-10 independent replicates. **(G)** Representative micrographs (of *pac1*Δ cells) and quantitation of spindle positioning phenotypes in the indicated yeast strains. Bars represent mean ± SD, and diamonds the fraction of mispositioned anaphase spindles in each replicate (n = 3-4 independent replicates).

To test the role of linker-Pac1 contacts in dynein-Pac1 binding, we engineered mutations (S1290A M1294A N1346A T1348A L1349A; see asterisks in Fig. 6F) into an otherwise wild-type GST-dynein_MOTOR_ (to test motility), and a monomeric dynein_MOTOR_^MT-U^ fragment that binds best to Pac1 monomers, and thus robustly forms 2 dynein_MOTOR_:1 Pac1^dimer^ complexes: that with both AAA1^WB^ and AAA3^WA^ mutations (Fig. 1). Single molecule motility assays reveal mutations at site^linker^ have no apparent effect on dynein motility (Fig. 7C). We next assessed the relative binding of the wild-type and mutant motors to monomeric and dimeric Pac1. Mutations at site^linker^ significantly reduces binding to Pac1^ΔN^, indicating that the Pac1-linker contacts indeed play a large role in governing high affinity binding at site^ring^ (Fig. 7D). We next sought to test our hypothesis that the Pac1-linker contact, and the resulting downward translation of Pac1 toward the stalk, occludes Pac1 binding at site^stalk^. If true, then assembly of 2 dynein_MOTOR_:1 Pac1^dimer^ complexes would be disrupted by the linker mutations since a Pac1^dimer^ would now occupy both site^ring^ and site^stalk^ instead of just site^ring^. This indeed is true, as the ability of the site^linker^ mutant to form these complexes was almost eliminated (Fig. 7E). In fact, in lieu of 2:1 complexes, the linker mutant assembles 1:1 complexes, indicating the linker-Pac1 contacts play no role in high affinity binding of the motor domain to Pac1 dimers.

If the linker-Pac1 contacts are responsible for the 1:1 binding mode, then a full-length AAA3^WA^ mutant with mutations at site^linker^ would assemble into 1:2 instead of 1:1 complexes with Pac1. This indeed proves to be the case (Fig. 7F). Finally, to test the functional importance of site^linker^, we engineered the mutations into an otherwise wild-type dynein locus in yeast cells, and assessed the position of the anaphase yeast. This reveals that mutations at site^linker^ are sufficient to disrupt dynein activity to identical levels of those cells lacking Pac1 (*pac1*Δ; Fig. 7G). Thus, site^linker^-Pac1 contacts, and assembly of 1:1 complexes, are critical for in-cell dynein function.

## Discussion

It is well established that various conditions modulate the affinity of dynein for Pac1 and Lis1. This includes whether dynein is bound or unbound from microtubules, as well as the particular nucleotide conditions^4, 42^. Although we recently uncovered the structural basis for microtubule binding affecting their affinity^4^, the precise role for nucleotides has been lacking. Here we show that multiple AAA domains within the dynein motor coordinate to affect not just Pac1 binding affinity, but also the stoichiometry of their binding. This includes AAA1, AAA3, and AAA4. We further find that dynein-Pac1 binding stoichiometry directly impacts whether Pac1 can relieve the autoinhibited phi conformation of dynein. Finally, cryo-EM analysis of full-length wild-type dynein reveals that the structural basis for the distinct binding modes lies in the direct interactions between the post-powerstroke linker and Pac1.

Negative stain EM analysis reveals that assembly of 1:1 full-length dynein:Pac1 complexes (in response to *e*.*g*., AAA1-ADP/AAA3-apo) results in the adoption of an apparent open conformation of dynein. In addition to being in an open state, the motor domains of the full-length dynein appear to be held in relatively stable orientations with respect to each other. Given the 2 dynein_MOTOR_:1 Pac1^dimer^ complexes sample a similarly limited inter-head distance, but exhibit a high degree of rotational movement (unlike Pac1 bound to full-length dynein), the stable configuration of the Pac1bound motor domains in the full-length complex is likely due to steric constraints imposed by both the Pac1 dimerization domain and the full-length dynein complex. Although the advantage or functional consequences of such a configuration are currently unclear, we hypothesize this conformation helps to promote assembly of dynein with dynactin and a cargo adaptor. This is in light of data indicating that phi-disrupting mutations strongly promote DDA complex assembly^2^. This conformation may also somehow assist dynein in binding microtubules (as has been observed^14, 31, 42, 53^), as the motor domains appear to be in a high microtubule-binding affinity state as predicted by the buttress-stalk conformation. Note that our data cannot distinguish between whether Pac1 opens dynein, or whether it simply stabilizes an open conformation.

Our cryo-EM data with full-length dynein shows that Pac1 stabilizes an apo-AAA3 pocket. Given the lack of dynein particles with an empty AAA3 pocket in the absence of Pac1, an apo-AAA3 is likely a short-lived state. Pac1 appears to be an opportunistic binder that strongly favors those dyneins that release ADP from AAA3 (possibly due to the post-2 state of the linker), and consequently binds and enriches for their presence. This suggests a mechanism by which Pac1 can open the phi state in cells: providing Pac1 is present at sufficiently high local concentrations, immediately subsequent to ADP release from AAA3, Pac1 binds due to its high affinity for site^ring^ and site^linker^ in this state. Although unclear, it is unlikely that ADP release from AAA3 is coordinated between the two motor domains within a dimer. Thus, once bound to one motor via site^ring^, the second WD40 within the Pac1 dimer is well placed to immediately bind the second motor domain after it releases ADP from its AAA3 domain.

Our findings show the importance of the linker in coordinating dynein-Pac1 binding stoichiometry. The importance of site^linker^ and the 1:1 complex it coordinates is highlighted by the fact that mutating this site results in a severe loss-of-function phenotype in cells (equivalent to loss of Pac1), in spite of this mutant having no measurable reduction in assembling the 1:2 complex (Fig. 7E and F). This region, which is the key effector of dynein’s powerstroke^45, 54-56^, has been proposed to be directly regulated by a bound Pac1^7, 51^. However, instead of Pac1 blocking the linker’s motion, as has been proposed^7^, we show that the post-2 linker position is required for high affinity 1:1 complex formation, and that the linker is also stabilized in a modified post-2 state by Pac1 binding. This contact may also be important for impacting AAA3 dynamics and stabilizing this pocket in an apo state. In light of the proposed importance of the post-2 linker in dynein stepping behavior^1, 52^, the Pac1-stabilized post-2* state and/or the apo-AAA3 pocket might account for the observed impacts of Pac1 and Lis1 on dynein force generation^42, 53^. The importance of the linker-Pac1 contact also explains why addition of ATP + Vi weakens affinity for Pac1 at site^ring^ (see AAA3^WA^ in Fig. 3A) and reduces 1:1 dynein:Pac1 binding stoichiometry for the AAA3^WA^ mutants (Figs. 1D, 4B, S1B). Treatment with ATP + Vi has been shown to enrich for a pre-powerstroke linker configuration^8, 54, 55, 57^. Such a bent linker would be expected to effectively eliminate site^linker^, and thus shift the binding from 1:1 to 1:2 stoichiometry, as we observe.

The linker position also likely accounts for results from our binding data with the monomeric dynein_MOTOR_ fragment (Fig. 1). Specifically, those conditions that enrich for the 2 dynein_MOTOR_ :1 Pac1^dimer^ species (*e*.*g*., apo-AAA3/ADP-AAA1) may do so because they promote a post-2 linker position. This is supported by our cryo-EM data (Fig. 6D), which show that ∼50% of the dynein motor domains in the minus-Pac1 conditions exhibit a post-2 linker state (*i*.*e*., are primed for Pac1 binding). Addition of Pac1 results in the elimination of dyneins with a post-2 linker (*i*.*e*., they are likely converted to a Pac1-bound motor with post-2*), but only a modest reduction in those with a post-1 linker (which are unbound from Pac1), likely due to the latter being in a low Pac1-binding affinity state.

Although we currently do not understand the functional significance of the 1:2 full-length dynein:Pac1 complex (in response to *e*.*g*., AAA1-ATP/AAA3-ATP), our negative stain images clearly show that this complex is not in an open state, but rather a ‘phi-like’ state. We speculate that this complex is structurally similar to a recently identified Lis1-bound phi-dynein^58^. As predicted by our binding assays, this complex possesses an ADP-AAA1 and ADP-AAA3. Unlike phi dynein, phi-Lis1 lacks C2 symmetry, which may explain our inability to obtain high quality 2D averages. We posit the following model to account for these distinct dynein-Lis1 complexes (Fig. 8): (1) Phi dynein, which has ADP bound at AAA1 and AAA3^1^ is primed to bind a Lis1 dimer via both site^ring^ and site^stalk^. (2) Dynein adopts a ‘phi-like’ conformation with either one or two Lis1 dimers bound. Given the high concentrations required to obtain two Lis1 dimers, it is more likely that dynein binds only one in the cell. (3) Dynein transiently adopts a partially open state, which results in ADP being released from AAA3, and the adoption of a post-2 linker state. (4) This transient conformation results in Lis1 dissociating from site^stalk^, and binding to site^ring^ on the 2^nd^ motor domain. (5) As a consequence of this new binding mode, Lis1 stabilizes dynein in a fully open conformation that is primed for assembly into motile DDA complexes. (6) Upon microtubule-binding and initiation of motility, Lis1 dissociates from the DDA complex^4^. Note that a recent high-resolution cryo-EM structure of a Lis1-bound DDA complex (with the Jip3 cargo adaptor) on microtubules showed a fully open/motile-competent configuration of dynein^5^. The microtubule-unbound dynein dimer in this complex is bound to 2 Lis1 dimers, with AAA1 and AAA3 being both bound to ADP, conditions that we find promotes the 2:1 complex, and disfavors 1:1 formation. Given this complex includes dynactin and a cargo adaptor, it likely reflects a later point in the activation mechanism. Future studies will be required to tease out how and whether each of these distinct dynein-Pac1/Lis1 complexes relate to each other, and their respective roles in the dynein activation process.

**Figure 8:**
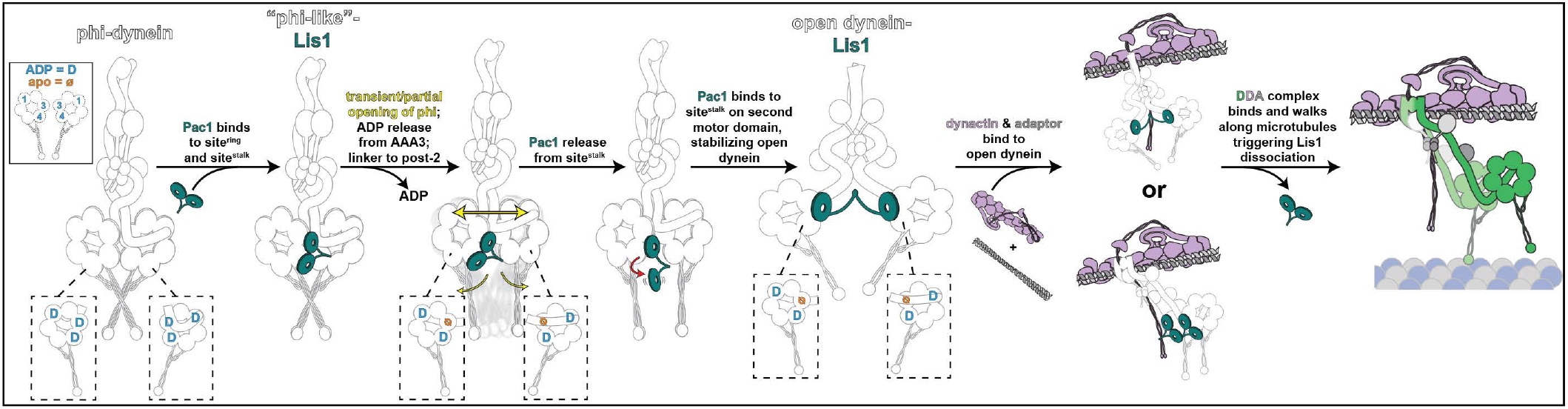
Model for Lis1 opening of phi dynein. We posit the following model to account for our data. Dynein primarily exists in the autoinhibited phi state with ADP bound to AAA1, 3 and 4^1, 2^. As a consequence of this nucleotide code, Lis1 can bind with one WD40 binding at site^ring^, and the other at site^stalk^. Our negative stain EM data indicate this binding mode does not result in dynein opening, consistent with a recent cryo-EM study^REF^. To release ADP from AAA3 and adopt a post-2 linker state, dynein partially/transiently opens, which consequently leads to Pac1 unbinding from site^stalk^, and binding to the 2^nd^ motor domain. This new binding mode stabilizes an open conformation of dynein, which more readily binds to dynactin and an adaptor. Although assembly of the dynein-dynactin-adaptor complex likely promotes adoption of a parallel arrangement of the motor domains^2^ (bottom), we cannot rule out dynein remaining in the Lis1-stabilized open conformation we observe by negative stain EM (top). Finally, upon binding to microtubules and initiating a motility event, Lis1 dissociates due to a weakened affinity at site^stalk^ (ref.^4^).

## Acknowledgements

We are grateful to members of the Markus, DeLuca, and Zhang labs for valuable discussions. Electron microscopy was done at the University of Colorado, Boulder EM Services Core Facility in the MCDB Department, with the technical assistance of facility staff. This work utilized the Alpine high performance computing resource at the University of Colorado Boulder. Alpine is jointly funded by the University of Colorado Boulder, the University of Colorado Anschutz, and Colorado State University. We would like to thank K. Zhou, J. Lin, M. Llaguno, and S. Wu, for their help with cryo-EM data collection at Yale Cryo-EM facility. The Yale Cryo-EM Resource is funded in part by the NIH grant S10OD023603. We thank J. Wang at National Cancer Institute for the help of cryo-EM data collection. This research was, in part, supported by the National Cancer Institute’s National Cryo-EM Facility at the Frederick National Laboratory for Cancer Research under contract 75N91019D00024. This work was funded by the NIH/NIGMS (R35GM139483 to S.M.M., and R35GM142959 to K.Z.).

## Author contributions

S.M.M. designed the study. I.C.G, B.R.I, W.D.T., A.H.I., S.C.G., S.V.B., and S.M.M. purified proteins for binding assays, and negative stain EM. I.C.G, B.R.I, W.D.T, S.C.G., and S.M.M. performed in vitro binding assays. J.S.B. and S.M.M. performed and analyzed live-cell microscopy for spindle positioning assays. P.C. and J.Y. acquired and analyzed cryo-EM data, and built PDB models with support from K.Z. P.C. and S.M.M. generated figures. S.M.M. wrote the manuscript. All authors edited and revised the manuscript. S.M.M. and K.Z. acquired funding.

## Materials and Methods

### Media and strain construction

Yeast strains are derived from W303 or YEF473A^59^ and are available upon request. We transformed yeast strains using the lithium acetate method^60^. Strains carrying mutations or tagged components were constructed by PCR product-mediated transformation^61^ or by transforming expression plasmids encoding for affinity-tagged proteins (see Tables S2 and S3). All mutagenesis was confirmed by whole plasmid sequencing. To integrate dynein_MOTOR_ expression plasmids, the plasmids were first digested with either ApaI or SbfI (both of which cut within the *URA3* gene; the latter was used for those plasmids with SRS^CC^) prior to transformation into a yeast strain deleted for the native *DYN1* gene. Strains overexpressing wild-type or mutant Pac1 were generated by transforming pRS306:GAL1p:8xHIS-ZZ-2xTEV-Pac1-FLAG-SNAPf (wild-type or mutant) linearized by digestion with ApaI. Integration of all plasmids were confirmed by diagnostic PCR. Yeast synthetic defined (SD) medium was obtained from Sunrise Science Products. (San Diego, CA).

### Plasmid and BACmid construction

A plasmid encoding for an affinity-tagged wild-type dynein_MOTOR_ fragment was generated as a starting point for the production of numerous mutants used throughout the manuscript (to enable simpler and more rapid engineering of mutants; see Table S2). This plasmid was made by amplifying the entire gene expression cassette from a yeast strain that encodes for this same fragment (SMY2675; see Table S3) using PCR primers that span from the 5’ end of the *GAL1p* promoter to the 3’ end of the HaloTag situated at the N-terminus of dynein_MOTOR_. This plasmid, which was assembled into pRS306 (with a *URA3* selection cassette) via traditional Gibson assembly, includes the following elements: GAL1p:ZZ-2xTEV-6His-dynein_MOTOR_-HaloTag (P1532). See Table S2 for mutations engineered into this plasmid via Gibson assembly.

For expression of biochemical quantities of high purity full-length yeast dynein complex, we generated a multicistronic plasmid that contains expression cassettes for all four dynein complex genes (*DYN2, DYN3, PAC11*, and *DYN1*; all codon-optimized for insect cells), with N-terminal affinity tags (6His-ZZ) followed by 2xTEV cleavage sites and a SNAPf tag on *DYN1*. This plasmid (P825) was assembled using the biGBac technique^62^. In brief, codon optimized open reading frames for *DYN2, DYN3* and *PAC11* were each assembled into pLib, whereas a codon optimized 6His-ZZ-2TEV-SNAPf-*DYN1* was assembled into pbiG1a (all of which contain 5’ polH promoters and 3’ SV40 terminators). Gene expression cassettes for *DYN2, DYN3* and *PAC11* (i.e., each with polH promoter and SV40 terminator) were amplified from their respective pLib plasmid, and then assembled into pbiG1b to generate pbiG1b:*DYN2/DYN3/PAC11*. The *DYN2/DYN3/PAC11* poly-gene cassette and 6His-ZZ-2TEV-SNAPf-*DYN1* were then combined together into pbiG2ab by cutting them out of their respective pbiG1 plasmids (by PmeI) and assembling them into PmeI-digested pbiG2ab, generating pbiG2ab: *DYN2/DYN3/PAC11/*6His-ZZ-2TEV-SNAPf-*DYN1* (hereafter referred to as pbiG2ab:yeast-dynein). Mutations indicated throughout the paper (*i*.*e*., K2424A, E2488Q, or the site^linker^ mutations) were engineered directly into this plasmid. All plasmids were validated by whole plasmid sequencing.

To generate BACmids (used for transfection into Sf9 cells), pbiG2ab:dynein-dynein (wild-type or mutant) were each transformed into DH10 EMBacY cells (Geneva Biotech), according to the manufacturer’s protocol. Proper transposition and BACmid generation was confirmed by blue/white screening. Resulting colonies were inoculated into LB media supplemented with 7 µg/ml gentamycin, 10 µg/ml tetracycline and 50 µg/ml kanamycin, grown overnight at 37°C, and purified as previously described^2, 33^. BACmids were stored at 4°C and used within 2 weeks for subsequent virus production (see below).

### Protein purification

Purification of dynein_MOTOR_ fragments (GAL1p:ZZ-2xTEV-6His-dynein_MOTOR_-Halo, GAL1p:ZZ-2TEV-6His-GFP-GST-dyn1-motor, or similar) was performed as previously described with minor modifications^23^. In brief, yeast cultures were grown in YPA medium supplemented with 2% galactose for 3-6 hours, collected, washed with cold water and then resuspended in a small volume of water. The resuspended cell pellet was drop-frozen into liquid nitrogen and then lysed in a coffee grinder. After lysis, 0.25 volumes of 4X dynein lysis buffer (1X buffer: 30 mM HEPES pH 7.2, 50 mM potassium acetate, 2 mM magnesium acetate, 0.2 mM EGTA) supplemented with 1 mM dithiothreitol (DTT), 0.1 mM Mg-ATP and 0.5 mM Pefabloc SC or protease inhibitor tablets (Pierce) (concentrations for the 1X buffer) was added, and the lysate was clarified by centrifugation at 310,000 x g for 1 hour. The supernatant was then incubated with IgG sepharose 6 fast flow resin (GE) for 1-3 hours at 4°C, which was subsequently washed three times in 5-10 ml lysis buffer, and twice in 5-10 ml TEV buffer (50 mM Tris pH 8.0, 150 mM potassium acetate, 2 mM magnesium acetate, 1 mM EGTA and 10% glycerol) supplemented with 0.005% Triton X-100, 1 mM DTT, 0.1 mM Mg-ATP and 0.5 mM Pefabloc SC. To label protein for single molecule assays (some GST-dynein_MOTOR_-HaloTag fragments), approximately 2-5 µM JFX650 Halo dye was added, and incubated for 10-20 minutes at room temperature, after which the resin was washed with 3 × 10 ml of TEV buffer. The resin was then incubated with TEV protease for 30 minutes at room temperature (for monomeric dynein_MOTOR_ fragments) or 1 hour at 16°C (for GST-dynein_MOTOR_-HaloTag fragments). After digestion with TEV, the bead-buffer slurry was filtered through an Ultrafree-MC microcentrifuge filters (to collect liquid phase) and aliquoted, flash-frozen in liquid nitrogen, and stored at -80°C. Protein quality was assessed by SDS-PAGE and mass photometry. Some dynein_MOTOR_ mutants were not sufficiently pure for mass photometry-based binding assays, and required a subsequent polishing step. To do so, the TEV-cleaved protein was diluted 10-20-fold in buffer A (20 mM Tris, pH 8.0, 1 mM magnesium acetate, 10% glycerol, 0.1 mM ATP, 1 mM DTT), and then injected on to a monoQ 10/100 GL pre-equilibrated in buffer A using an AKTA Pure. After injection, the column was washed with 2-5 column volumes of buffer A, and then bound proteins were eluted using 20 column volumes of a linear gradient from buffer A to buffer B (20 mM Tris, pH 8.0, 1 mM magnesium acetate, 1 M NaCl, 10% glycerol, 0.1 mM ATP, 1 mM DTT). The peak fractions (typically between 25-35% buffer B) with the highest purity (determined by mass photometry) were pooled, concentrated, aliquoted, flash-frozen and then stored at -80°C. Note that one mutant in particular – dynein_MOTOR_^MT-U/K2766A^ – required a gel filtration step instead of ion exchange to obtain protein of sufficient purity for our mass photometry-based binding assays. To this end, the concentrated peak fractions from the monoQ were applied to a Superose 6 10/300 using GF150 (25 mM HEPES pH 7.4, 150 mM KCl, 1 mM MgCl_2_, 1 mM DTT, 0.1 mM Mg-ATP), and the highest purity fractions (determined by mass photometry) were then pooled and concentrated before aliquoting, freezing, and storage at -80°C.

GST-dimerized dynein motors (expressed from yeast cells expressing ZZ-2xTEV-6His-GFP-GST-dynein_MOTOR_-HALO) used for mass photometry experiments were tandem affinity purified from yeast cell cultures as previously described^63^. Briefly, following cell lysis, 0.25 volumes of 4X NiNTA dynein lysis buffer (1X buffer: 30mM HEPES pH 7.2, 150 mM potassium acetate, 2 mM magnesium acetate and 10% glycerol) supplemented with 1 mM beta-mercaptoethanol, 0.1 mM Mg-ATP and 0.5 mM Pefabloc SC (concentrations for the 1X buffer) was added, and the lysate was clarified as described above. The supernatant was then bound to NiNTA agarose for 1 h at 4°C, which was subsequently washed three times in 5 ml NiNTA lysis buffer. The protein was eluted in NiNTA lysis buffer supplemented with 250 mM imidazole by incubation on ice for 20 min. The eluate was then diluted with an equal volume of 1X dynein lysis buffer (see above), which was then incubated with IgG Sepharose 6 fast flow resin for 1 h at 4°C. The beads were washed three times with dynein lysis buffer, and then twice with TEV buffer and the protein was eluted as described above (TEV protease treatment for 30 min at room temperature). Eluted protein was applied to a size-exclusion resin (Superose 6; Cytiva) preequilibrated in TEV buffer using an AKTA Pure system. Peak fractions (determined by absorbance at 260 nm and SDS–PAGE) were pooled, concentrated, aliquoted, flash-frozen and then stored at −80 °C.

Purification of ZZ-TEV-Pac1-SNAPf was performed as previously described^23, 63^. Briefly, protein was prepared as described above for dynein_MOTOR_ fragments with the following two differences. Instead of incubating with TEV protease for 30 minutes at room temperature, TEV digest was performed at 4°C overnight. Instead of polishing the protein by anion exchange chromatography, the TEV eluate was applied to a size exclusion chromatography resin (Superose 6, Cytiva) that was equilibrated in TEV buffer supplemented with 1 mM DTT using an AKTA Pure. Peak fractions (determined by absorbance at 260 nm and SDS-PAGE) were pooled, concentrated, aliquoted, flash-frozen and then stored at -80°C.

The intact full-length yeast dynein complex was expressed in and purified from insect cells (Sf9 cells) using a similar protocol to that used for the human dynein complex^64^. Briefly, a 6-well dish was seeded with 2 ml Sf9 cells at 5 × 10^5^ cells/ml, which were maintained in SF 900 II SFM Medium (Life Technologies). Cells were transfected with 2-6 µg of BACmid DNA using FuGENE HD transfection reagent, and incubated at 27°C in a sealed box with a damp towel inside. The efficiency of transfection and virus production was monitored via YFP fluorescence. 4-7 days following transfection (when 95-100% cells were expressing YFP reporter), the virus (P1) was collected by transferring media to a sterile tube. The virus was amplified by adding 1 ml of P1 virus to 50 ml Sf9 cells at 5 × 10^5^ cells/ml, which were incubated at 27°C in a flask shaking at 140 rpm. The resulting P2 virus was collected by centrifuging cell suspension, and transferring virus-containing media to a sterile tube, which was maintained at 4°C. For protein production, 20 ml of P2 was used to infect 2 L of Sf9 cells at 1.5 × 10^6^ cells/ml. 66-70 hours after infection, the cells were harvested (2000 x g, 20 min), washed with lysis buffer (50 mM HEPES, pH 7.4, 100 mM NaCl, 10% glycerol, pH 7.2), pelleted again (1810 x g, 20 min), and resuspended in an equal volume of lysis buffer supplemented with 1 mM DTT, 0.1 mM Mg-ATP, and 1 mM PMSF. The resulting cell suspension was drop frozen in liquid nitrogen and stored at −80°C. For protein purification, ∼30 ml of fresh dynein lysis buffer supplemented with cOmplete protease inhibitor cocktail (Roche), 1 mM DTT, and 0.1 mM Mg-ATP was added to the frozen cell pellet, which was then rapidly thawed in a 37°C water bath prior to incubation on ice. Cells were lysed in a dounce-type tissue grinder (Wheaton) using ∼60 strokes. Subsequent to clarification at 310,000 x g for 1 hour at 4°C, the supernatant was applied to 3-4 ml of IgG sepharose fast flow resin (Cytiva) pre-equilibrated in lysis buffer, and incubated at 4°C for 3-5 hours. Beads were then washed with 30-50 ml of lysis buffer, and then 20-30 ml of TEV buffer (50 mM Tris pH 7.4, 150 mM potassium acetate, 2 mM magnesium acetate, 1 mM EGTA, 10% glycerol, 1 mM DTT, 0.1 mM Mg-ATP). The beads were then incubated with 300 µg of TEV protease overnight at 4°C. The next morning, the recovered supernatant was applied to either a Superose 6 gel filtration column (Cytiva) or a TSKgel G4000 column. Peak fractions were pooled, concentrated, aliquoted, flash frozen, then stored at -80°C. Sample quality was assessed using mass photometry, SDS-PAGE, and/or negative-stain electron microscopy prior to cryo-EM grid preparation.

### Negative Stain Electron Microscopy

EM grids were prepared by applying freshly purified dynein complexes to glow discharged carbon coated 200 mesh copper grids. After a ∼1 minute incubation, 2% uranyl acetate was added. Micrographs were collected on a FEI Tecnai F20 200kV TEM equipped with a Gatan US4000 CCD (model 984), at a nominal magnification of 90,000X with the digital pixel size 1.449 angstroms. All image analysis was performed in Relion 4.0 on the University of Colorado Boulder High Performance Computer Cluster, Alpine. Particles were manually picked from ∼20-30 micrographs (∼300-400 particles), which were used to generate low resolution 2D class averages. These 2D averages were then used to autopick particles used to generate our final 2D averages.

### Mass Photometry

All proteins were diluted to either 50 nM, or a fraction greater or lesser thereof (as indicated by X-value in Figures 4 and S1) in assay buffer (50 mM Tris, pH 8.0, 150 mM potassium acetate, 2 mM magnesium acetate, 1 mM DTT) with or without added nucleotide (1 mM of each). 1-2 µl of each protein was then mixed 1:1 (to 25 nM of each), incubated for 1-2 minutes, and then diluted 1:5 on the stage (2 µl of mixed protein + 8 µl of same buffer with or without nucleotide) to a final concentration of 5 nM immediately prior to image acquisition. For apo conditions, residual ATP from the protein preparation was depleted using apyrase by mixing 4.5 µl of respective dynein_MOTOR_ protein with a 0.5 µl of apyrase (NEB), and incubating for 30 minutes at room temperature. 1-minute movies were acquired using Refeyn MP, and all images were processed and analyzed using Discover MP. Calibration was performed with beta-amylase and thyroglobulin.

### Cryo-EM sample preparation and data collection

Dynein concentration was diluted to 2 mg/ml (from 3 mg/ml) in GF150 buffer with or without Pac1 and incubated on ice for 1 h before cryo-EM sample vitrification (Fig. S2B). 3 μL sample was applied to Quantifoil holy carbon grids (R2/1, 300 mesh gold or R2/1, 400 mesh) and incubated in the chamber of the Vitrobot Mark IV unit (FEI) for 5 seconds at 4 °C and 100% humidity. Subsequently, the grids were blotted for 3-5 seconds and plunged into liquid ethane. The grids were then screened, and data were collected on a 200 keV Glacios electron microscope (Thermo Fisher Scientific) with a K3 direct detection camera (Gatan) with a magnification of 45000x, physical pixel size of 0.868 Å and total dose of 40 e-/Å^2^. In total, 2, 179 (for dynein alone) and 7,036 (for dynein-pac1 complex) movies at a defocus from -1.2 μm to -2.7 μm were acquired and data collection was automated using SerialEM^65, 66^.

### Cryo-EM image processing

Preprocessing steps, including motion correction, CTF estimation, and particle picking, were conducted either in cryoSPARC Live^67^ or via an inhouse script utilizing MotionCor2^68^, GCTF^69^, and Gautomatch. Scripts for real-time data transfer and on-the-fly preprocessing are available for download at https://github.com/JackZhang-Lab. The following reconstruction steps were conducted in cryoSPARC^67^ (Fig. S3C and D).

For both datasets, the initial particles were picked using a blob picker or template matching and extracted at a box size of 128 at 3.47 Å pixel size (bin 4). The particles were subjected to several rounds of 2D classification to obtain high-quality class averages of dynein motor domain. Ab initio reconstruction followed by heterogeneous refinement of these particles (6 classes) reveal medium resolution (5-8 Å) of the motor domains. Manual inspection was performed for different classes and similar classes were merged. Two different classes were identified, and the particles were re-extracted at a box size of 284 at 1.157 Å pixel size for high-resolution reconstruction. Each class was subjected to homogeneous refinement and two rounds of global CTF, local CTF and local refinement. The resolutions of all maps were estimated by Fourier shell correlation calculations^70, 71^ embedded in cryoSPARC. Local resolution analysis^72^ was performed in cryoSPARC.

### Model building and refinement

The previous published yeast dynein motor domain structures (PDB: 4AKI^73^, 4W8F^57^, 7MGM^3^) were used as initial models. A manual fitting step was conducted in Chimerax^74^ to fit individual AAA domain to the cryo-EM maps. Namdinator^75^ was then employed to automatically refine the model into the cryo-EM map.

All models underwent iterative refinement using Phenix real-space refinement 1.21rc1_5190^76^ and manual rebuilding in COOT^77, 78^. The quality of the refined models was assessed using MolProbity^79^ integrated into Phenix, with statistics reported in Table S1.

### Plots and molecular graphics

Plots were generated using GraphPad Prism. Molecular graphics were prepared using UCSF ChimeraX^74^.

## Supplementary Information

**Table S1:** Cryo-EM Data Collection, Refinement, and Validation Statistics.

|  | Yeast full-length dynein alone in 0.1mM ATP condition |  | Yeast full-length dynein with Pac1 in 0.1mM ATP condition |  |
| --- | --- | --- | --- | --- |
| Description | Class-1:<br>Apo-AAA1,<br>ADP-AAA3,<br>post-1 | Class-2:<br>ADP-AAA1,<br>ADP-AAA3,<br>post-2 | Class-1:<br>Apo-AAA1,<br>ADP-AAA3,<br>post-1 | Class-2:<br>ADP-AAA1,<br>Apo-AAA3,<br>Pac1-bound,<br>post-2 |
| Deposition Code | PDB: XXXX<br>EMDB: XXXX | PDB: XXXX<br>EMDB: XXXX | PDB: XXXX<br>EMDB: XXXX | PDB: XXXX<br>EMDB: XXXX |
| Facility | Yale ScienceHill-Cryo-EM facility |  |  |  |
| Microscope | Glacios |  |  |  |
| Voltage (kV) | 200 |  |  |  |
| Camera | K3 |  |  |  |
| Magnification | 45k |  |  |  |
| Pixel Size (Å) | 0.432 (super resolution) |  |  |  |
| Total Electron Exposure (e-/Å²) | 40 |  |  |  |
| Defocus Range (µm) | 1.5-2.7 |  |  |  |
| Num of mics | 2,169 |  | 7,036 |  |
| Initial Particles | 1,190,329 |  | 2,521,271 |  |
| Final Particles | 63,405 | 53,499 | 93,983 | 178,195 |
| Symmetry Imposed | C1 | C1 | C1 | C1 |
| Initial models | 4AKI | 4W8F | 4AKI | 4W8F<br>7MGM |
| Map pixel size | 1.302 | 1.302 | 1.157 | 1.157 |
| Map Resolution (Å) (FSC 0.143) | 3.3 | 4.1 | 3.2 | 3.3 |
| Map sharpening B-factor (Å²) | 83 | 64 | 68 | 57 |
| Model Composition |  |  |  |  |
| Non-hydrogen atoms | 21,020 | 20,038 | 21,020 | 23902 |
| Protein residues | 2,584 | 2,934 | 2,584 | 2,940 |
| Ligands | MG:1<br>ATP:1<br>ADP:2 | ATP:1<br>ADP:3 | MG:1<br>ATP:3<br>ADP:2 | MG:2<br>ATP:1<br>ADP:2 |
| Model vs. Data |  |  |  |  |
| FSC Map to Model (Å) (FSC 0.5) | 3.0 | 4.5 | 3.5 | 3.5 |
| Correlation coefficient (mask) | 0.77 | 0.85 | 0.83 | 0.83 |
| B factors (Å²) |  |  |  |  |
| Protein | 49.53 | 184.81 | 49.53 | 63.85 |
| Ligand | 21.00 | 136.06 | 21.00 | 41.41 |
| R.m.s deviation |  |  |  |  |
| Bond length (Å) | 0.005 | 0.003 | 0.004 | 0.004 |
| Bond angles (°) | 0.952 | 0.663 | 0.582 | 0.608 |
| Validation |  |  |  |  |
| Molprobrity score | 1.79 | 2.08 | 1.79 | 1.81 |
| Clashscore | 10.09 | 17.34 | 10.12 | 10.01 |
| Rotamer outliers (%) | 0.04 | 0.00 | 0.04 | 0.04 |
| Ramachandran plot |  |  |  |  |
| Outliers (%) | 0.00 | 0.08 | 0.00 | 0.1 |
| Allowed (%) | 3.91 | 4.84 | 3.91 | 4.12 |
| Favored (%) | 96.09 | 95.08 | 96.09 | 95.77 |
| Rama-Z (whole) | 0.4 | 0.16 | 0.4 | 0.69 |

**Figure S1:**
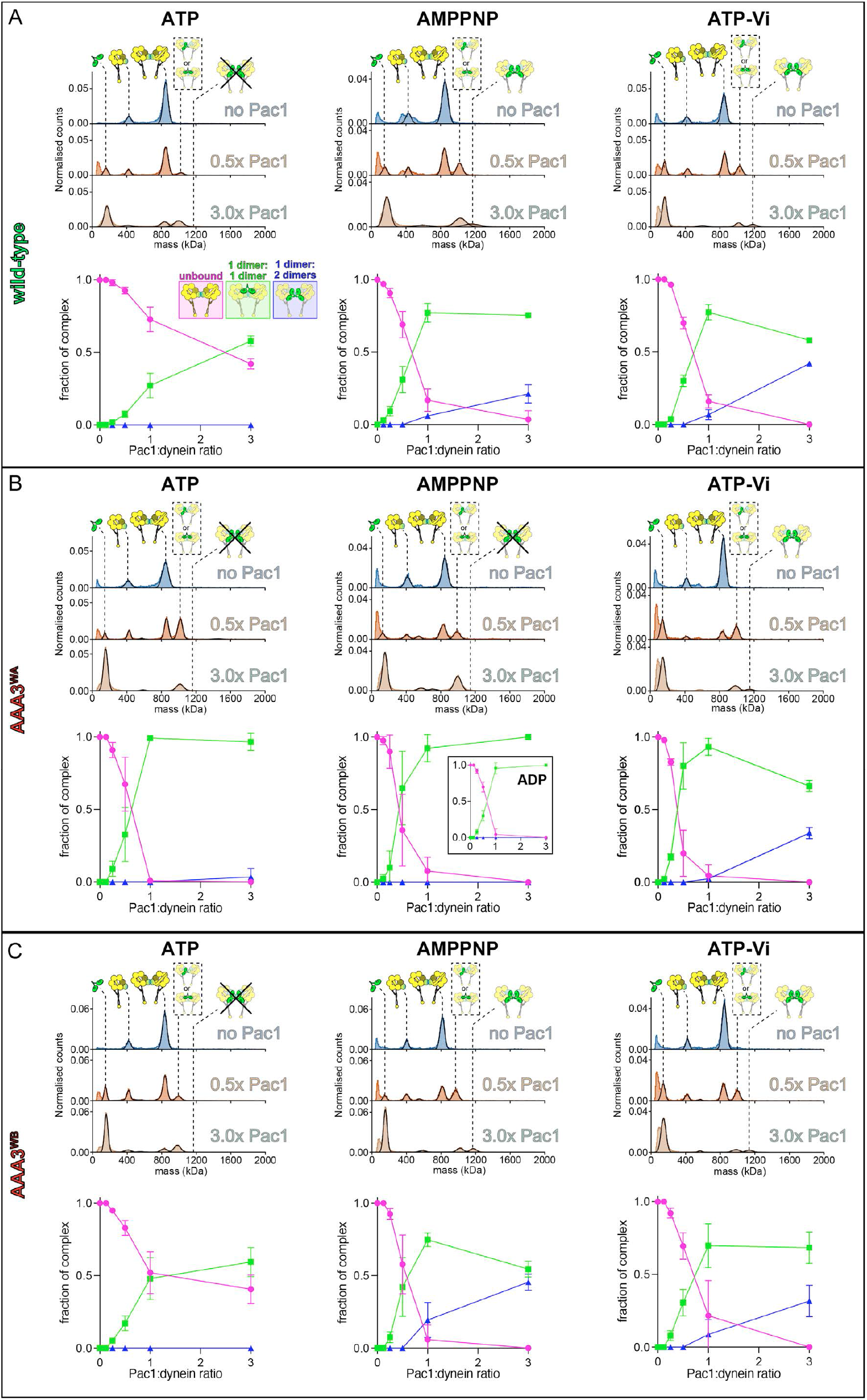
Nucleotide occupancy coordinates stoichiometry of binding between Pac1 and GST-dimerized dynein_MOTOR_. Representative mass photometry histograms and associated plots (mean fraction of binding ± SD; n = 3 independent replicates for each) depicting extent and stoichiometry of binding between Pac1 and indicated GST-dynein_MOTOR_ at a range of dynein:Pac1 ratios.

**Figure S2:**
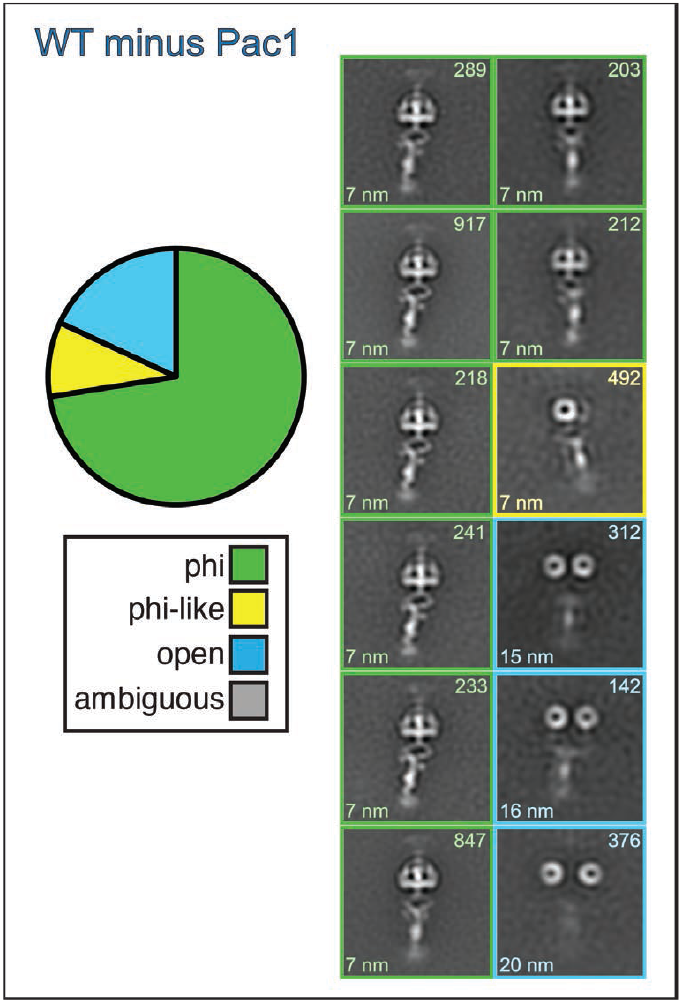
Negative stain EM of wild-type full-length dynein. Representative 2D class averages showing indicated full-length dynein (purified from Sf9 cells) in the absence of Pac1. Pie graph shows relative fraction of indicated conformational state (determined from 2D averages). Numbers indicate the number of particles in each class (top) and the measured distance between the motor domains (from center-to-center; bottom).

**Figure S3:**
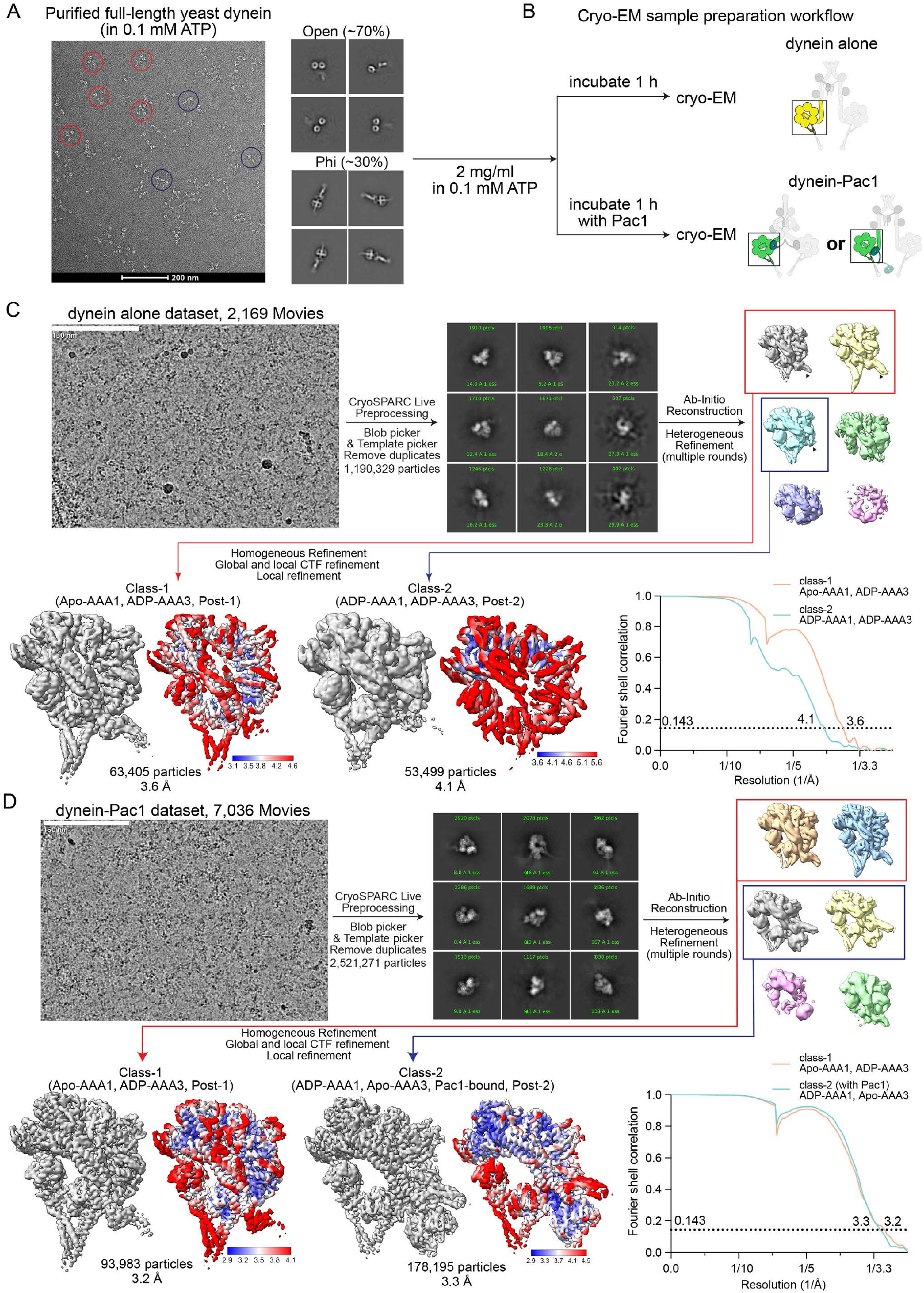
Cryo-EM data processing workflow. **(A)** A representative negative staining image of purified yeast full-length dynein with corresponding 2D class averages. Red and blue circles indicate the open and phi dynein, respectively. **(B)** Cryo-EM sample preparation workflow of dynein alone or in complex of Pac in 0.1 mM ATP concentration. **(C and D)** Cryo-EM image processing of the dynein alone dataset **(C)** or the dynein-pac1 dataset **(D)**. Representative 2D class averages of motor domains are shown. Cryo-EM maps, local-resolution analysis and FSC plots are shown.

**Figure S4:**
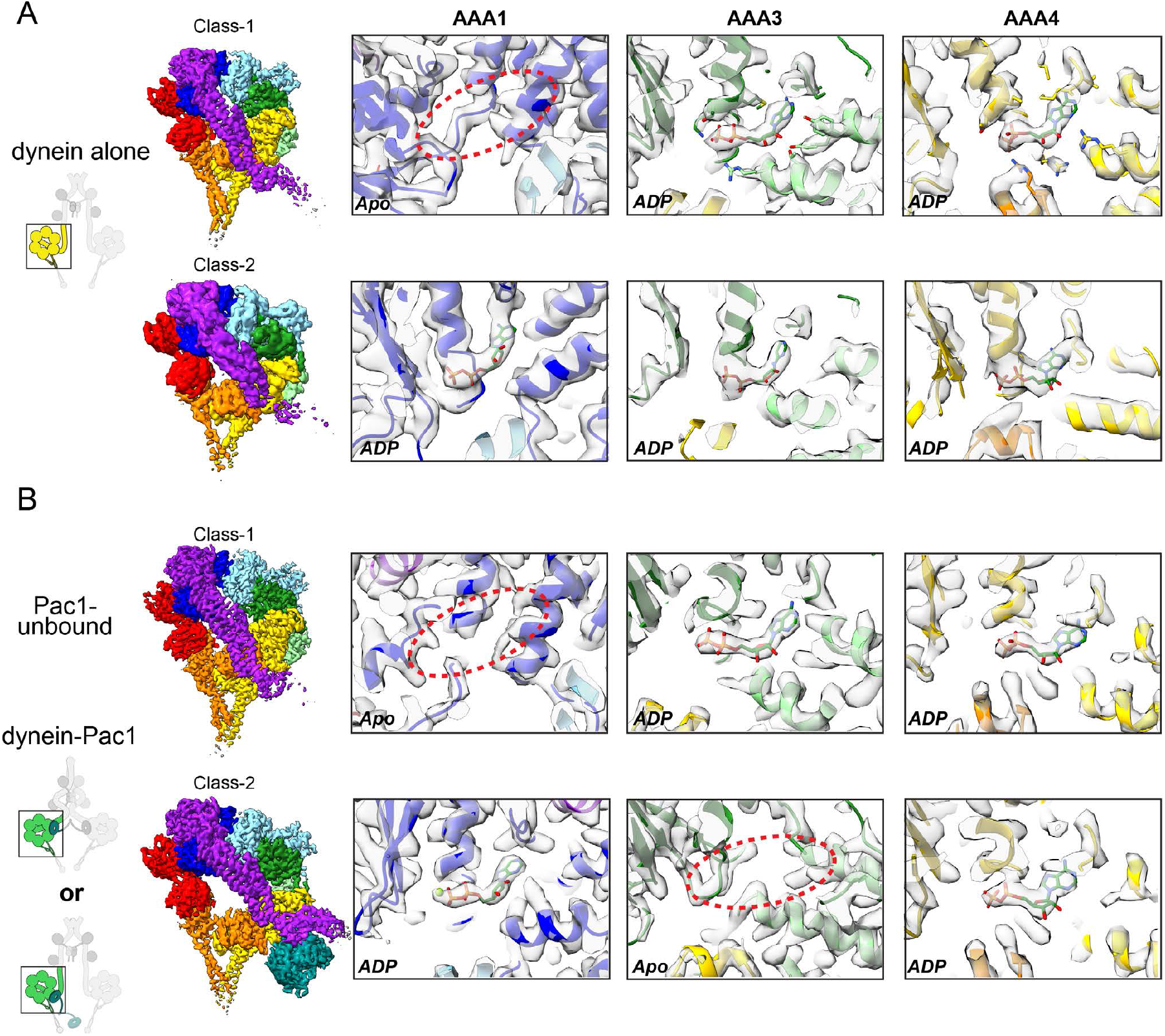
Nucleotide states of AAA1, AAA3 and AAA4 from each cryo-EM map. **(A and B)** Two classes from the dynein alone dataset **(A)** or from the dynein-pac1 dataset **(B)**. The Cryo-EM maps are shown in transparent surface mode fitted with PDB models. The red dashed circles highlight the missing nucleotide in the pocket (apo state).

